# Residue-specific dominant-negative mutant of ubiquitin reveals functional selectivity as Ubp14 deubiquitinase inhibitor

**DOI:** 10.64898/2026.05.18.725830

**Authors:** Amrita Arpita Padhy, Saanya Yadav, Subhashree Sahoo, Kummari Shivani, Lahari Reddy Balireddygari, Himanshu Joshi, Parul Mishra

## Abstract

Multifunctional proteins encode specificity through nuanced molecular interactions that operate in the presence of abundant wild-type molecules. Previous studies on multifunctional proteins have shown how certain regulated interactions parse complex biological activities into discrete functional modules. Discrete interaction edges can function as regulatory nodes whose perturbations selectively remodel proteostasis output, especially under disease conditions. However, whether such interaction specificity can be harnessed to selectively manipulate functional modules and reveal residue-level control principles, remains largely unexplored. Here, we screened for dominant negative mutations that impact partner-selective functions linked to Leu8, a critical binding residue on ubiquitin. Our results reveal that the Leu8Ala mutation specifically leads to accumulation of polyubiquitinated proteins and decreased levels of free ubiquitin suggesting loss of deubiquitinase function in yeast cells. Cellular and biochemical analyses establish that the Leu8Ala variant of ubiquitin specifically inhibits the yeast deubiquitinase, Ubp14, and its human homolog, USP5. We further demonstrate remodelling of the binding interface with increased interface contacts for the variant and Ubp14 complex. The variant shows a higher inhibitory potency compared to the wild-type ubiquitin and can inhibit Ubp14 both as unconjugated and as conjugated ubiquitin chain indicating the strength of its inhibitory function. Our results provide mechanistic insight into how edge perturbations in ubiquitin can reveal critical nodes that impact selective functions and thus fine-tune the cellular proteostasis network for therapeutic benefit.

## Introduction

Ubiquitin is a core protein of cellular proteostasis being a versatile post translational modifier that regulates functions of protein localization, trafficking, protein-protein interactions and protein degradation in cells (Bett, 2016; Finley, 2009; Yau and Rape, 2016). Ubiquitin associates with almost all cellular proteins via an enzymatic cascade involving ubiquitin activating (E1), conjugating (E2s) and ligating (E3s) enzymes. The free pools of ubiquitin are maintained in the cells by the deubiquitinases (DUBs) that function to remove the ubiquitin moiety from its substrates (Clague et al., 2015; Komander and Rape, 2012). Although the repertoire of E2s, E3s, and DUBs has expanded considerably with cellular complexity, ubiquitin exhibits high level of sequence conservation across species (Kaminskaya et al., 2024). Within its broad interactome, ubiquitin is the shared ligand in its molecular complexes formed with E2s, E3s, and DUBs suggesting that the structural properties of ubiquitin might impose stringent constraints to govern the specificity of its interactions with the diverse proteins in the cell. Studies show that interaction interfaces are closely clustered on the surface of ubiquitin implying multiple and overlapping binding functions of individual residues of ubiquitin (Mavor et al., 2016; Michel et al., 2024; Roscoe et al., 2013a; Sloper-Mould et al., 2001a). Hence, changes in single amino acids can potentially bias the interaction equilibria selectively towards one or more complexes in the ubiquitin network within the cells. Although the molecular mechanisms of ubiquitin centric pathways are continuously being explored, it is imperative to understand the residue level contributions that regulate the diversity of functions and how subtle perturbations to ubiquitin structure can balance proteostasis outcome.

Structurally complex proteins like Hsp90 and calmodulin discriminate among their diverse clientele of >500 proteins in the cell despite of lacking a dedicated binding site. Their multi-domain architecture allows them to undergo large conformational shifts (>10Å) from apo to ligand bound forms to sample between discrete pool of interactors (Babu et al., 1985; Lopez et al., 2021; Tidow and Nissen, 2013). Contrarily, ubiquitin and ubiquitin family of proteins display a simple and evolutionarily conserved compact beta grasp fold with a well- defined surface and hydrophobic core regions where structural mobility is largely restricted to its C-terminus and loop regions (Hochstrasser, 2009; Perica and Chothia, 2010; Van Der Veen and Ploegh, 2012; Vijay-kumar et al., 1987). Consequently, ubiquitin exhibits restrictive and localized conformational dynamics in these flexible regions that populate solution structures recognised by selective DUBs, E2s or E3s (Lindorff-Larsen et al., 2005). Interestingly, the minor population of conformationally selected states shows minimal changes in the overall structure of ubiquitin but follow remodelling of the UIM/DUBs-Ub interface to enhance the affinity of its interactions (Beriashvili et al., 2024; Lange et al., 2008; Wlodarski and Zagrovic, 2009). These studies imply that large conformational rearrangements are limited in ubiquitin and molecular properties of its surface residues might impose a specificity switch to regulate ubiquitin signalling in the dynamic cellular environment.

Dominant negative mutants are exceptionally powerful tools to study essential genes, multimeric proteins and signalling pathways that cannot be targeted by the conventional gene deletion strategies (Gerasimavicius et al., 2022; Herskowitz, 1987; Marutani et al., 1999; Meyer-Schuman et al., 2023; Miller et al., 1988; Schweisguth, 1999). Studies employing overexpression of dominant negative mutants that sequester critical factors have revealed rate limiting steps in complex biological pathways or inform about the spatio-temporal dynamics of functions (Gerasimavicius et al., 2022; Padhy et al., 2023). Global perturbations in cellular functions have been widely observed with expression of dominant negative mutants for multimeric proteins (Papa et al., 2014). For example, dominant negative mutation even in one subunit of the DNA binding domain of p53, reduces the transcriptional activation efficiency of the p53 mixed tetramer appreciably reducing the downstream expression of many DNA damage response proteins (De Vries et al., 2002). Dominant mutants have also been helpful to identify critical structural transitions for tissue specific selectivity of interactors. For example, mutations in yeast mitochondrial Hsc4p that dominantly locked the chaperone in ATP bound state demonstrated tissue specific difference in the clientele of this chaperone (Elefant and Palter, 1999). Dominant negative mutants of ubiquitin like I44A, G76A and K48R utilized to probe pathway specific effects have demonstrated global disruption of ubiquitination (Finley et al., 1994; Terrell et al., 1998). Although these nodal studies have dissected ubiquitin conjugation (G76A Ub traps E2-Ub conjugates), linkage specificity (K48R Ub disrupts proteasome degradation pathway) and recognition (I44A inhibits recognition by all UBDs), selective dominant interference with specific interactors of ubiquitin remains unexplored.

DUBs are positioned for a unique regulatory role in the ubiquitin network where they remodel the ubiquitin chain topologies to decide the fate and function of cellular substrates (Mevissen and Komander, 2017; Reyes-Turcu et al., 2009a). They govern the flux of proteasomal and non-proteasomal pathways by regulating the steady state levels of free and conjugated pools of ubiquitin (Trulsson et al., 2022). While the E2s interact with ubiquitin via its C-terminal tail conjugation and via appropriate docking of the hydrophobic patch finally incorporating E3 into this adduct, DUBs recognize the precise conformations of ubiquitin as their conjugated forms or with the substrates. Consequently, even subtle structural perturbations in ubiquitin are capable of impairing DUBs-Ub association with measurable shifts in conjugated and free ubiquitin in cells. This heightened sensitivity to conformational and geometrical changes in ubiquitin allows DUBs to be an ideal system to uncover residue level determinants of functional selectivity in ubiquitin.

This study uncovers residue and substitution specific precise functional interference in cellular ubiquitination pathways. In our previous work we have demonstrated that numerous dominant variants exist across ubiquitin protein which can expose critical functional nodes in the ubiquitin network. We advanced this concept by resolving dominant interference to the level of single residue and its specific substitution identifying leucine 8 as a uniquely sensitive surface residue of ubiquitin. We demonstrated that the L8A mutant ubiquitin dominantly and selectively inhibits a yeast deubiquitinase, Ubp14, revealing an unanticipated residue-level determinant of DUBs function. Biochemical assays showed enhanced binding of this variant to Ubp14 and its human homolog, the USP5 which caused the deubiquitinase to be quenched from the cellular pool thereby disturbing the ubiquitin homeostasis. Molecular simulations revealed that the interaction is stabilized through local interface effects without significantly perturbing the ubiquitin’s overall fold. While in vivo studies demonstrated that the inhibitory effects of the mutant ubiquitin is primarily via its conjugated form, the in vitro activity analysis showed that Ubp14 preferentially associated with the unconjugated mutant ubiquitin. Our study demonstrated a strategy to utilize rationally designed ubiquitin mutants as molecular probes to investigate the complex ubiquitin signalling and uncover hidden regulatory dependencies suitable for therapeutic targeting.

## Results

### Structure guided analysis of residue specific dominant negative substitutions in ubiquitin reveal distinct ubiquitination profiles

Ubiquitin consists of several well-defined hydrophobic patches that play a crucial role in mediating interactions with ubiquitin-binding proteins in the cell (Fig 1A). The most common interaction site is the I44-patch, comprising of L8, I44, and V70 residues that bind to the helical ubiquitin-interacting motif (UIM), ubiquitin-associated helices (UBA) or zinc finger containing domains of numerous ubiquitin binding proteins (Randles and Walters, 2012). The I36 patch engages with a few DUBs and E3 ligases, while the F4 patch and the TEK box are shown to regulate receptor endocytosis and mitosis respectively (Haas and Wilkinson, 2008; Sloper-Mould et al., 2001a). While the specificities towards these diverse interactions might be regulated via conformational dynamics in each of the hydrophobic patches, we focussed on the I44 patch which should be most sensitive to structural perturbations, being the most interaction-dense surface among other patches. We further reasoned that a residue residing in the interface core of I44 patch with E1, E2, E3 or DUBs would contribute strongly to the ubiquitin-enzyme interactions than a residue at peripheral positions. Hence, we systematically evaluated the extent to which L8, I44 and V70 amino acids of this patch engaged with these enzyme classes by comprehensively surveying all the co-crystal structures of ubiquitin with E1, E2, E3 and DUBs (Fig 1B and 1C). We first collected 168 PDB structures representing all the available structures of wild type ubiquitin in complex with E2s, E3s, DUBs and other non-enzymatic proteins. We then filtered out the structures that represented the same ubiquitin-enzyme complexes obtained by alternative methods to avoid over-representation of any complex. We also excluded all ternary complexes from the dataset to prevent confounding interpretations arising due to secondary contacts within these complexes. We finally retained only 41 high resolution binary complexes of ubiquitin with E2, E3 and DUBs to access the primary interaction interfaces. Although, the surveyed structures included complexes from diverse species but since ubiquitin is highly conserved across eukaryotes, our structural analysis likely reflected conserved structural principles of interactions rather than species-specific characteristics. We then quantified the solvent-accessible surface area (SASA) for L8, I44 and V70 in each unique ubiquitin-enzyme complex using Pymol (Fig 1D and Supplemental table 1). Residue-specific buried surface area was calculated as the difference between the SASA value of each residue in the isolated ubiquitin structure (PDB ID: UBQ1) and its SASA value in the ubiquitin-enzyme complex structure. Comparison of the fraction buried area for L8, I44 and V70 across enzyme classes revealed a differential engagement of the I44 patch residues. The E2s associate variably with the three residues of the I44 patch with no emerging residue preference for being most buried across all the structures analysed suggesting diverse binding geometries across E2s. The V70 residue showed to be relatively more buried in the ubiquitin: E3 ligases structures. Interestingly, the L8 residue of ubiquitin showed a high buried surface area and tight clustering of this property in all the ubiquitin: DUBs structures compared to its location in ubiquitin: E2 or E3 complexes suggesting that L8 is a core residue of ubiquitin and DUBs interface (Fig 1D). This analysis demonstrated the presence of residue level asymmetry in how E2s, E3s and DUBs engage with the hydrophobic patch residues of ubiquitin and provided a structural basis for investigating if substitutions at L8 position could selectively bias DUB interactions and generate class specific dominant interference. We have previously observed that a larger number of chemically diverse dominant negative mutations exist at the L8 position in ubiquitin compared to I44 and V70 positions (Padhy et al. 2023). In contrast to other hydrophobic patches (F4, I36 and TEK box) which displayed ∼45% of the substitutions to be dominant negative, the I44 patch showed ∼ 25% of dominant negative substitutions with most of them localized at the L8 residue of the patch (Fig S1A). Hence, to probe for subtle substitutions that would generate dominant effects in the global interaction network of ubiquitin, we focussed on the L8 amino acid of the I44 hydrophobic patch. We selected all the substitutions at the L8 position of ubiquitin which showed dominant negative phenotype in our bulk fitness analysis screen and examined the growth defects and levels of ubiquitinated proteins when each mutant was individually overexpressed in the W303 yeast strain. We observed a range of toxic growth phenotype for the mutant proteins (Fig 1E). Ubiquitin with L8A, L8D, L8H, and L8Q mutation showed a moderate toxicity (∼35 %), while L8M, L8R, L8Y, L8E, L8G, L8S and L8P mutant ubiquitin had almost 50-60% reduced growth with L8N being the most toxic mutant compared to cells overexpressing wild-type ubiquitin along with endogenous ubiquitin expression. Each ubiquitin mutant was overexpressed under the galactose promoter which led to ∼3.5 fold overexpression as estimated by the levels of the wild-type free ubiquitin under these expression conditions (Fig S1B). We observed that the substitutions of leucine to non-polar and hydrophobic residues like methionine and tyrosine conferred a higher toxicity compared to the structurally different amino acids like aspartate and alanine suggesting that dominant negative phenotypic effects are based on more complex parameters of interface geometry. To assess if any of these mutations are strongly destabilizing for ubiquitin structure, we also analysed the ΔG values of each mutant protein using FoldX. All the substitutions at the L8 position showed ΔG values < +1Kcal/mol suggesting that they are not destabilizing mutations (Supplemental table 2). To investigate the changes in the levels of conjugated and free ubiquitin associated with the toxic growth phenotypes, we performed western blot analysis of yeast cells overexpressing the selected L8 mutants for 3 hrs in galactose-containing medium. Unexpectedly, we observed a reduced accumulation of ubiquitinated proteins in most of the L8 mutants compared to the control cells overexpressing the wild-type ubiquitin (Ub^WT^). Interestingly, the L8A mutant ubiquitin (Ub^L8A^) showed the highest level of polyubiquitinated proteins and lowest levels of free ubiquitin in the yeast cells compared to other mutations at the same position (Fig 1F). This distinct ubiquitination profile is consistent with impaired deubiquitinase activity in cells (Park and Ryu, 2014; Trulsson et al., 2022), suggesting that the Ub^L8A^ competitively interferes with deubiquitnase function of a cellular DUBs. Notably, reducing the galactose concentration to 0.5%, partially rescued the growth defect that was observed upon Ub^L8A^ expression in the yeast cells confirming that the observed phenotype was a dose dependent effect directly attributable to the Ub^L8A^ expression (Fig S1C). Together, these findings demonstrated that in the I44 hydrophobic patch, L8 residue plays a critical role for the ubiquitin: DUBs interactions and specific substitutions at this position can dominantly interfere with DUBs functions in cells.

**Figure 1:**
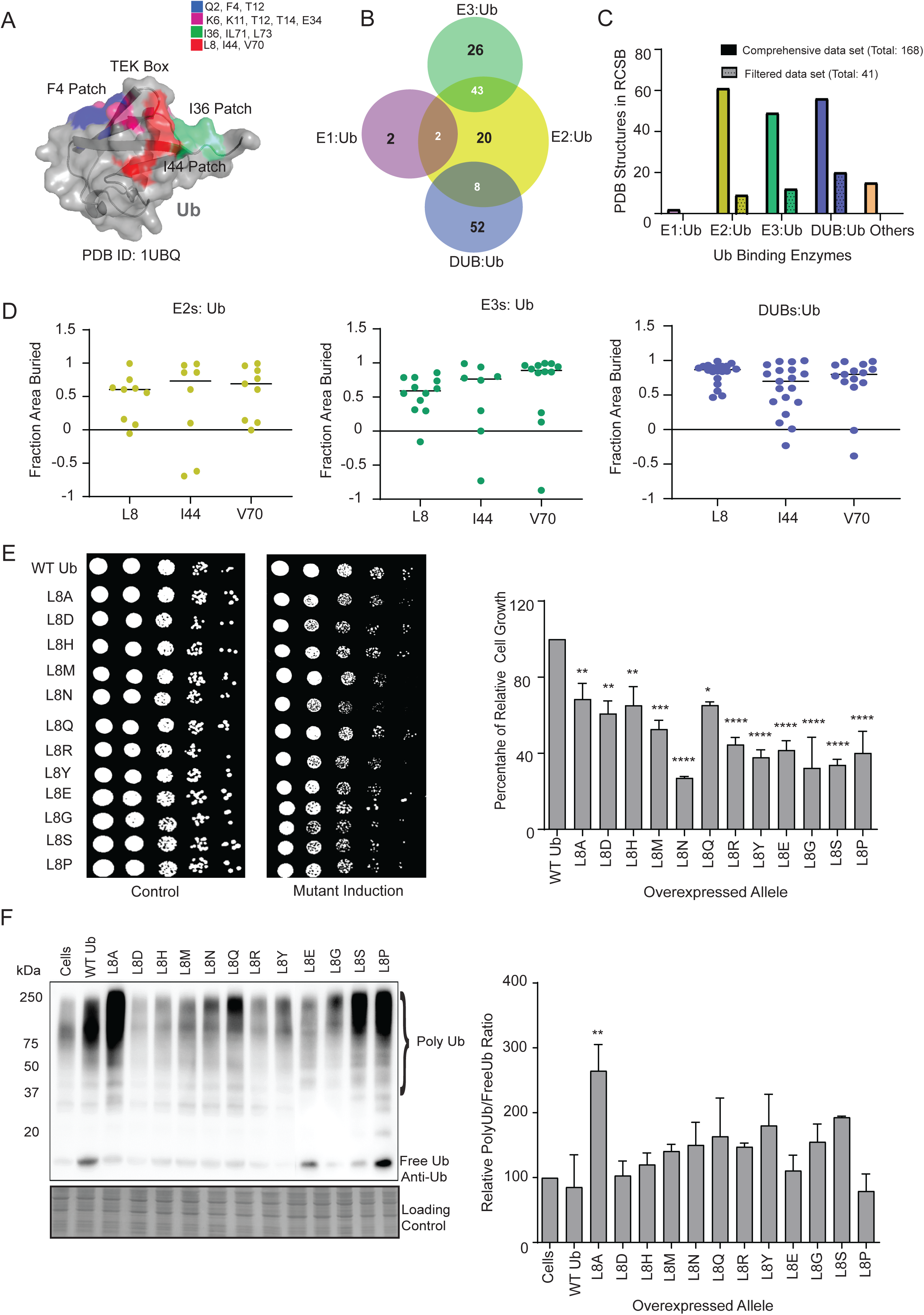
Structure guided analysis of residue specific dominant negative substitutions in ubiquitin reveal distinct ubiquitination profiles. (A). Cartoon representation of ubiquitin (PDB ID: 1UBQ) highlighting surface patches involved in protein–protein interactions. The F4, TEK box, I36, and I44 patches are shown in blue, pink, green, and red, respectively. (B). Venn diagram showing the number of ubiquitin-bound structures in each enzyme category, with overlapping regions indicating structures where ubiquitin interacts with more than one enzyme partner. (C). Bar graph showing the total number of ubiquitin co-crystal structures available in the PDB database and the subset reviewed for further analysis. The complete list of PDB IDs is provided in Supplementary Table 3. (D). Fractional buried surface area plots for residues in the I44 patch grouped by enzyme class (E2s, E3s, and DUBs). Structures analyzed correspond to the curated set of unique ubiquitin co-crystal structures from panel. Solvent-accessible surface area (SASA) for each I44 patch residue was calculated using PyMOL. (E). Growth analysis of yeast cells overexpressing individual L8 mutants alongside an endogenous copy of WT Ub. Spot dilution assays were performed on tryptophan-deficient synthetic media, with dextrose as the control medium and galactose for mutant induction. The bar graph shows growth rates of ubiquitin variants relative to overexpressed WT Ub in cells grown on solid medium for 2–3 days. Data are presented as mean ± SEM (n = 3 independent experiments). Statistical significance was determined using one-way ANOVA with Dunnett’s post hoc test comparing each mutant with wild type. (F). Western blot showing accumulation of polyubiquitinated proteins in cells expressing dominant-negative L8 ubiquitin mutants. Immunoblots were probed for ubiquitin, and the ratio of polyubiquitin to monoubiquitin (PolyUb/MonoUb) was quantified by densitometry. The bar graph represents quantification of polyubiquitinated species. Data are presented as mean ± SEM (n = 3 independent experiments). Statistical significance was determined using one-way ANOVA with Dunnett’s post hoc test comparing each mutant with wild type.

### L8A mutant ubiquitin dominantly inhibits Ubp14

Considering that ubiquitin deconjugation follows its enzymatic conjugation to substrates, we first sought to determine if Ub^L8A^ is competent for being incorporated into the polyubiquitin chains. We deleted the C-terminal glycine (G^75^G^76^) in the mutant sequence which is necessary for the enzymatic transfer of ubiquitin to substrates. We observed that the formation of high molecular weight species was abolished in Ub^L8AΔGG^ expressing cells similar to that observed for Ub^WTΔGG^ (Fig 2A). The Ub^L8AΔGG^ expressing cell also showed growth phenotype similar to wild type cells (Fig S2A). This suggested that the Ub^L8A^ is conjugated into the polyubiquitin chains and the toxic growth effect observed in cells expressing the mutant was not due to the free mutant form but due to its conjugated form. To confirm if the Ub^L8A^ upon expression, comprised a significant component in the accumulated polyubiquitin chains, we expressed the HA tag version of Ub^WT^ and Ub^L8A^ in the W303 yeast strain and analysed the polyubiquitinated species for HA by immunoblotting. We observed HA tagged high molecular weight species for Ub^L8A^ expressing cells suggesting that the mutant ubiquitin is efficiently incorporated into the polyubiquitin chains along with the endogenous ubiquitin forming mixed chains in the cells (Fig S2B). It has been previously shown that purified L8A/I44A double mutant of ubiquitin has a slower rate of E2 activation, but whether L8A mutant ubiquitin alone is deficient for forming polyubiquitin chains in the cells has not be tested. We expressed Ub^L8A^ and wild-type ubiquitin from a constitutive GPD promoter in the SUB328 yeast strain, where the endogenous ubiquitin is galactose induced. Western blot analysis of the cells grown in glucose containing yeast growth medium demonstrated the accumulation of polyubiquitinated proteins for both Ub^WT^ and Ub^L8A^, although the levels of polyubiquitinated proteins in Ub^L8A^ were reduced compared to the control cells (Fig S2C). Of note, we could only identify the polyubiquitinated species in early time points of growth in Ub^L8A^ expressing cells as this mutant is highly deleterious for cell growth when it is the only copy of cellular ubiquitin (Fig S2D). This suggested that leucine-8-alanine substitution alone in ubiquitin is structurally competent to form homotypic polyubiquitin chains even when the availability of wild-type ubiquitin is limited although the mutant is severely deficient in other cellular functions. Interestingly, these results demonstrated that Ub^L8A^ which strongly compromises all the cellular functions of ubiquitin as a single copy modulates some specific function/s of ubiquitin in cells in the presence of wild-type ubiquitin.

**Figure 2:**
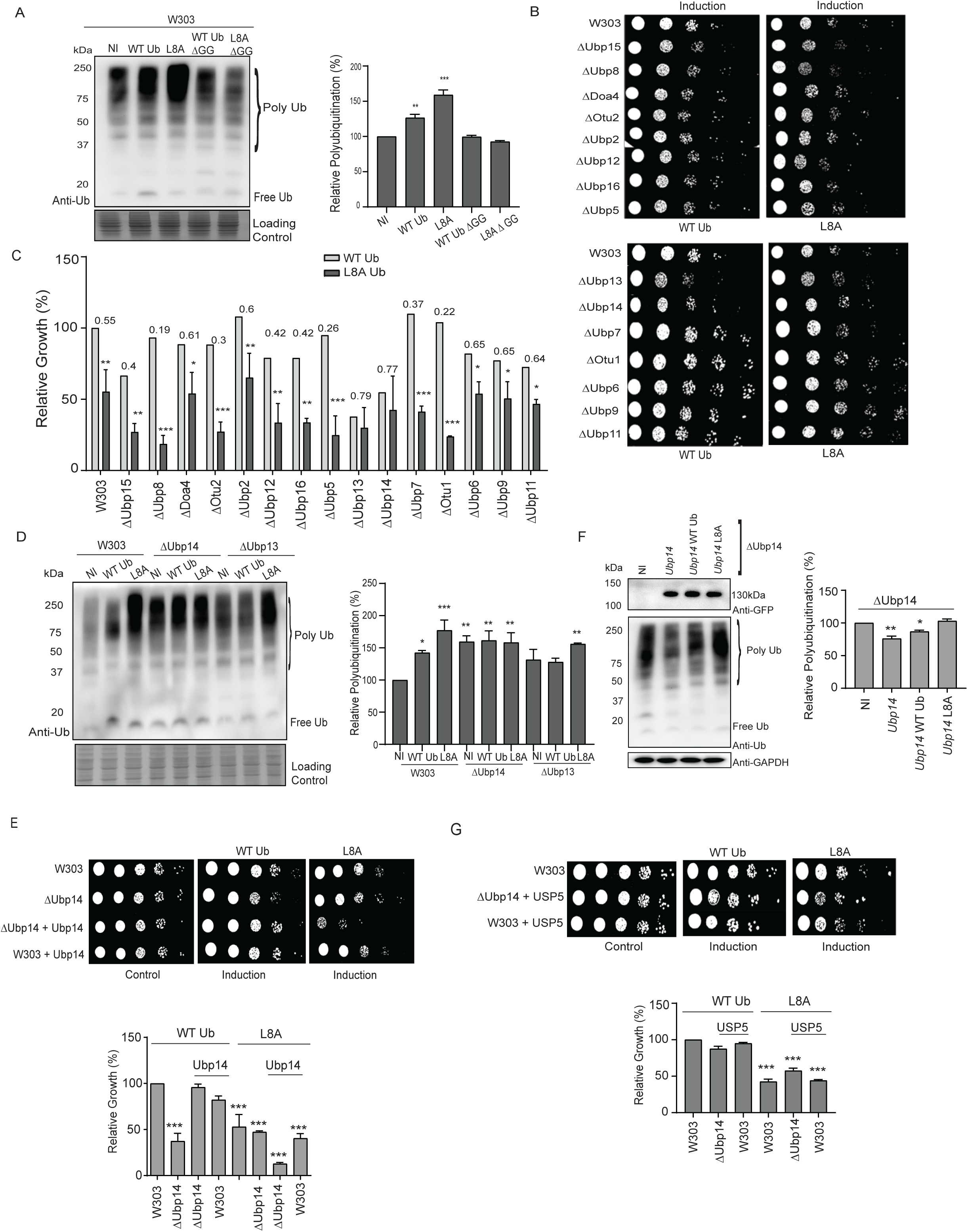
L8A mutant ubiquitin dominantly inhibits Ubp14. (A). Western blot and growth analysis of W303 cells expressing wild-type and ΔGG ubiquitin mutants grown on tryptophan-deficient synthetic media. Cells were cultured in dextrose (control) or galactose (mutant induction) for 6 hours. (B). Growth analysis of DUB deletion yeast strains expressing the ubiquitin mutant compared with strains expressing WT Ub. Spot dilution assays were performed on histidine-deficient synthetic media, with dextrose as the control medium and galactose for mutant induction. (C). Bar graph showing the fold change in growth of the mutant relative to wild type in each deletion strain grown on solid medium for 2–3 days. (D). Western blot showing accumulation of polyubiquitinated proteins in cells expressing mutant or WT Ub in the indicated DUB deletion strains. The bar graph represents quantification of polyubiquitinated species. Data are presented as mean ± SEM (n = 3 independent experiments). Statistical significance was determined using one-way ANOVA with Dunnett’s post hoc test comparing each mutant with W303. (E). Growth analysis of DUB deletion yeast strains with or without ectopic expression of Ubp14 in the presence of the ubiquitin mutant compared with WT Ub. Spot dilution assays were performed on histidine–uracil-deficient synthetic media, with dextrose as the control medium and galactose plus copper for induction of the mutant and DUB expression respectively. Bar graphs show quantification of growth on induction plates. Data are presented as mean ± SEM (n = 3 independent experiments). Statistical significance was determined using one-way ANOVA with Dunnett’s post hoc test comparing each mutant with wild type. (F). Western blot showing accumulation of polyubiquitinated proteins in DUB deletion yeast strains with or without ectopic expression of Ubp14 or USP5 in cells expressing the WT Ub and L8A. The bar graph represents quantification of polyubiquitinated species. Western blot of GFP tagged Ubp14 expression in Ubp14 overexpression samples and GAPDH as loading control are also shown. Data are presented as mean ± SEM (n = 3 independent experiments). Statistical significance was determined using one-way ANOVA with Dunnett’s post hoc test comparing each mutant with W303 wild type. (G). Growth analysis of DUB deletion yeast strains with or without ectopic expression of USP5 in the presence of the ubiquitin mutant compared with WT Ub. Spot dilution assays were performed on histidine–uracil-deficient synthetic media, with dextrose as the control medium and galactose plus copper for induction of the mutant and DUB expression respectively. Bar graphs show quantification of growth on induction plates. Data are presented as mean ± SEM (n = 3 independent experiments). Statistical significance was determined using one-way ANOVA with Dunnett’s post hoc test comparing each mutant with wild type.

To examine if the dominant negative effect of Ub^L8A^ is due to the inhibition of DUBs present in the yeast cells, we assessed the yeast growth across 15 individual yeast DUB deletion strains each ectopically expressing Ub^L8A^ (Fig 2B). We evaluated the fold change in growth of the cells overexpressing Ub^L8A^ relative to the cells overexpressing Ub^WT^ in the same genetic background (Fig 2C). Of note, deletion of some non-essential DUBs like ΔUbp6, ΔUbp7, ΔUbp9, ΔUbp11 and ΔOtu1 showed growth advantage while deletion of other non-essential DUBs tested in our panel had growth comparable to wild-type W303 yeast strain. ΔUbp3 showed a sick phenotype compared to W303 strain suggesting its critical role in cellular functions (Fig S2E). The growth of these DUBs deletion strains remained unchanged when Ub^WT^ was overexpressed in these strains (Fig 2B and 2C). Upon Ub^L8A^ overexpression, all the deletion strains except ΔUbp14 and ΔUbp13 exhibited growth disadvantage when compared to the same deletion strain with only Ub^WT^ overexpression. A moderate growth defect (∼45%) was observed in ΔUbp6, ΔUbp9, ΔUbp11, ΔDoa4, and ΔUbp2 strains while a stronger growth defect of ∼60% was seen in the remaining eight DUB deletion strains, indicating synthetic impairment of ubiquitin homeostasis in these DUBs deletion strains rather than a direct impairment of their function by Ub^L8A^ overexpression. Interestingly, Ub^L8A^ expressing ΔUbp14 and ΔUbp13 strains grew almost similar to their control cells suggesting preferentially interference of Ub^L8A^ with these DUBs. We also evaluated the growth effects of Ub^L8S^, a mutant that showed polyubiquitination profiles nearly similar Ub^L8A^ in the W303 cell, in the ΔUbp14 strain. Notably, expression of Ub^L8S^ showed an enhanced growth defect when expressed in the ΔUbp14 strain suggesting that it does not directly or primarily impact Ubp14 function (Fig S2F).

If Ub^L8A^ interferes with Ubp14 or Ubp13 functions specifically then we should observe a loss of accumulation of polyubiquitinated proteins in the deletion strain of these DUBs expressing Ub^L8A^. Hence, we examined the polyubiquitin levels in the ΔUbp14 and ΔUbp13 strains following induction of the wild-type and the mutant ubiquitin. We observed that there was no polyubiquitin accumulation in ΔUbp14 strain expressing Ub^L8A^, but, an increase in polyubiquitinated species was seen in ΔUbp13 strain expressing Ub^L8A^ (Fig 2D). These results suggested that as Ubp14 function are directly perturbed by Ub^L8A^ expression, hence its deletion suppressed both the toxic phenotypic effects and polyubiquitin accumulation by the mutant. In contrast, Ubp13 is not required for Ub^L8A^ induced chain accumulation but it might regulate the ubiquitination of critical substrates hence coupling the Ub^L8A^ expression mediated ubiquitin chain accumulation to cellular toxicity.

To validate if Ub^L8A^ specifically and dominantly alters the functions of Ubp14, we overexpressed Ubp14 ectopically in both W303 and the ΔUbp14 strain also expressing Ub^L8A^ and examined the growth effects on solid yeast growth media. Following the induction of both Ub^L8A^ (or Ub^WT^ in control cells) and Ubp14 in these strains, we observed a clear growth defect in ΔUbp14 cells compared to the control cells (Fig 2E). This toxic growth phenotype was also supported by the accumulation of polyubiquitnated proteins (Figure 2F) suggesting that Ub^L8A^ expression interferes with the ectopically expressed Ubp14. However, in the W303 strain, overexpression of Ubp14 did not further enhance the Ub^L8A^ mediated growth defects suggesting that Ub^L8A^ expression might be limiting compared to the levels of Ubp14 under the overexpression conditions. Hence when we decreased the Ub^L8A^ expression by inducing it in lower galactose concentration, we observed that the toxic growth phenotype of Ub^L8A^ was now recovered with overexpression of Ubp14 in these cells (Fig S2G). Together, these findings provided strong functional evidence that Ub^L8A^ acts as a selective inhibitor of Ubp14 function by trapping this deubiquitinase in the cells rather than broadly perturbing multiple deubiquitinases.

We next asked if Ub^L8A^ could also interfere with the cellular functions of USP5, the human orthologue of Ubp14 when expressed in yeast cells. To assess the feasibility of cross-species DUBs inhibition, we first examined the structural conservation between Ubp14 and USP5 in the ubiquitin binding domain. While the structure of human USP5 in complex with ubiquitin was available (PDB ID: 3IHP), we generated the yeast Ubp14: ubiquitin complex using the AlphaFold 3 server. Structural alignment revealed strong homology of the ubiquitin-binding interface in the two complexes, comprising of the catalytic cysteine and the histidine boxes (Fig S2H). Of note, the L8-interacting residues in USP5 aligned closely with the corresponding L8-binding residues in Ubp14, which suggested a conserved mechanism of ubiquitin recognition and potential cross-species specificity of the Ub^L8A^ towards USP5. To functionally test this possibility, we overexpressed the human USP5 gene in the ΔUbp14 strain co-expressing Ub^L8A^ and analysed the growth effects on solid yeast growth media. We observed a growth defect in this strain similar to that seen upon Ubp14 expression suggesting that Ub^L8A^ inhibited the function of USP5 similar to Ubp14 (Fig 2G) indicating functional complementation and conservation of DUB activity between yeast Ubp14 and human USP5. Together, these findings indicated that Ub^L8A^ selectively perturbed the cellular functions of Ubp14 and its human homolog USP5, highlighting the cross-species functional relevance of this mutant ubiquitin.

### L8A mutant ubiquitin impairs cell cycle progression and proteasomal degradation of substrates

Ubp14 is the only deubiquitinase in yeast which is known to disassemble unanchored polyubiquitin chains and replenish the free ubiquitin in cells (Amerik et al., 1997; Suresh et al., 2020). This unique function of Ubp14 strongly supports its critical role in maintaining the ubiquitin homeostasis and cellular health. As Ubp14 deletion is yeast cells is associated with impaired proteasome function, we examined if Ub^L8A^ mediated inhibition of Ubp14 reduced the proteasomal degradation of cellular proteins. For this, we measured the stability of two well-known ubiquitin-fusion degradation reporter proteins, Ub–Pro–β-galactosidase and Ub–Leu–β-galactosidase which are rapidly degraded via the proteasome. We transformed these genes into the W303 strain expressing either Ub^WT^ or Ub^L8A^ and monitored the steady state levels of these substrates using the cycloheximide pulse chase assay (Fig 3A). Compared to the control cells, Ub^L8A^ expressing cells showed a marked stabilization of both the substrates. The Ub-Pro-β-gal had a half-life of 15 min in the W303 cells but was not seen to be degraded even after 15 minutes when Ub^L8A^ was expressed (Fig 3A) and was stable till 30 min of the experiment (data not shown). A more pronounced stabilization was observed for Ub–Leu–β-gal from 5 minutes in the control cells to 15 minutes in cells having the mutant expression. This significant stabilization of the proteasome-targeted substrates would also be observed if the mutant impaired substrate recognition or substrate delivery to the proteasome impairing proteasome mediated degradation besides interfering with the Ubp14 deubiquitinase function. We ruled out any parallel direct effects of the mutant on the proteasome because the expression of Ub^L8A^ in a ΔUbp14 strain did not demonstrate any increase in the levels of polyubiquitnated proteins as compared to the ΔUbp14 strain alone (Fig 1D). This suggested that the observed defects in proteasomal degradation of substrates resulted from the direct inhibition of Ubp14 by Ub^L8A^ and disruption of the Ubp14 mediated ubiquitin homeostasis rather than from the intrinsic impairment of proteasome function by Ub^L8A^.

**Figure 3:**
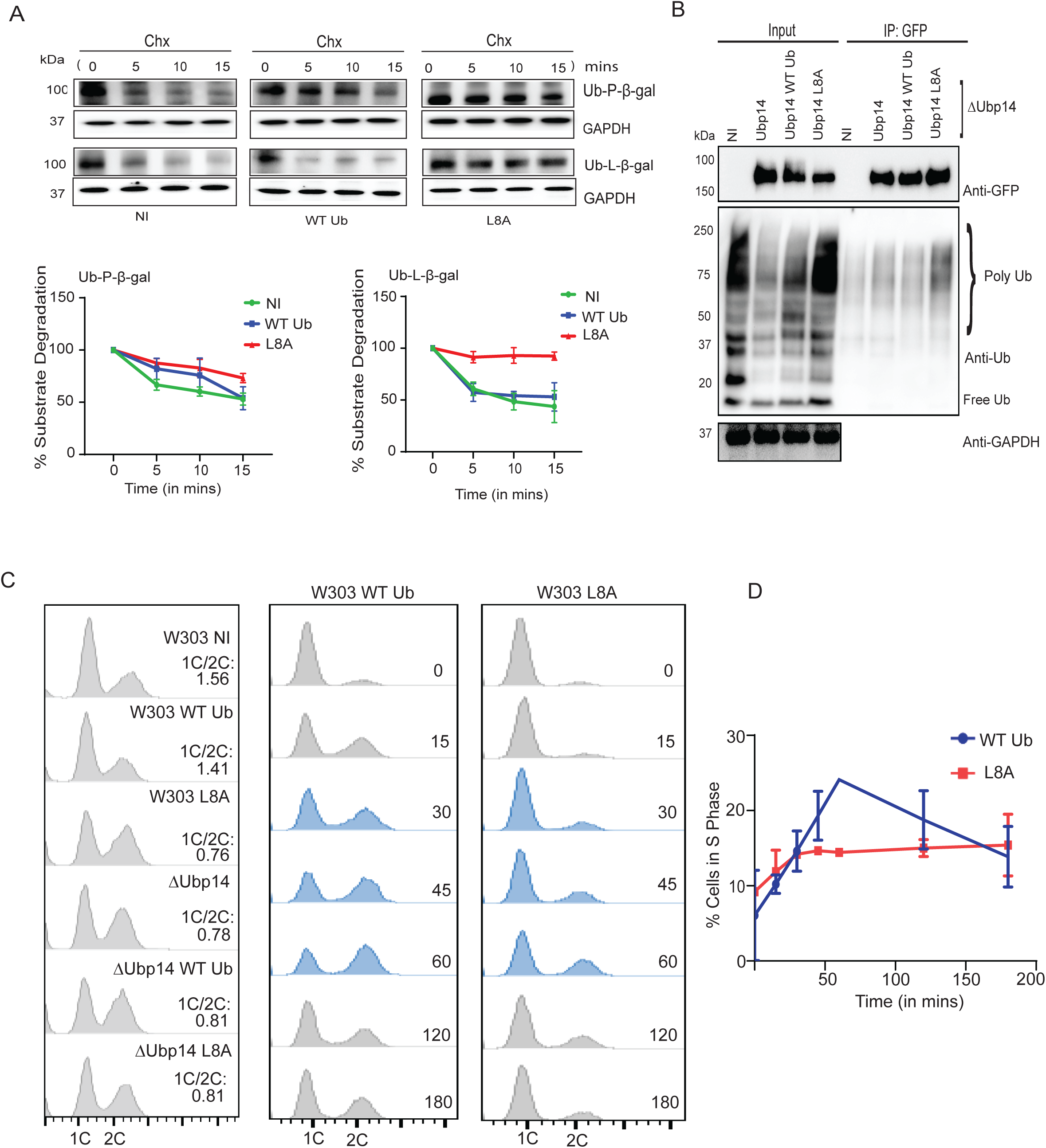
The L8A mutant ubiquitin impairs cell cycle progression and proteosomal degradation of substrates in a Ubp14-dependent manner. (A). Cycloheximide chase assays showing the half-life of the Ub–L–β-gal substrate in the indicated strains. Strains expressing the substrate were grown in uracil-deficient dextrose medium and induced in galactose medium for 6 hours. The lower panel shows quantification of cycloheximide chase assays from panel. Data represent the average of n = 3 experiments, with error bars indicating SEM. (B). Co-immunoprecipitation assay to assess ubiquitin levels in DUB-expressing cells with or without ubiquitin mutant induction. Samples were collected after 9 hours of copper and galactose induction for DUB and ubiquitin expression, respectively. Cell lysates were subjected to immunoprecipitation using an anti-GFP antibody and probed with the indicated antibodies. (C). FACS profiles showing DNA content of asynchronous W303 and ΔUbp14 cells grown in synthetic medium overexpressing WT Ub or L8A. The 1C:2C ratio is shown to highlight the decrease in the 1C population upon DUB inhibition or deletion. (D). Delayed entry into S phase in W303 L8A cells. W303 wild-type and L8A cells grown in synthetic medium were arrested in G1 with alpha factor and released into the cell cycle. Representative FACS plots are shown, with S-phase time points highlighted in blue. The percentage of S-phase progression for W303 wild-type and L8A cells is plotted as the average of n = 3 experiments, with error bars representing standard deviation.

Efficient degradation of ubiquitin-tagged substrates, requires efficient polyubiquitin chain binding and processing by the Ubp14. We next examined if functional interference of Ubp14 by Ub^L8A^ expression was due to the altered binding of Ubp14 to the polyubiquitin chains. We performed co-immunoprecipitation in yeast cells co-expressing GFP-tagged Ubp14 and Ub^L8A^ in ΔUbp14. We observed that Ubp14 was associated with polyubiquitinated proteins in cells expressing Ub^L8A^ or Ub^WT^ however, the Ub^L8A^ expressing cells showed a marked increase in levels of precipitated polyubiquitinated species (Fig 3B). This suggested that Ubp14 is not binding deficient, instead, it is strong association to polyubiquitin chains in vivo might impair its ability to disassemble the associated ubiquitin chains into free ubiquitin. This result highlights a “substrate trapping” effect of Ub^L8A^ expression presenting a proteostasis bottleneck of maintaining adequate levels of free and conjugated ubiquitin in the cells.

As Ub^L8A^ expression stabilized the reporter proteins, we hypothesised that this mutant ubiquitin could also affect the turnover of key cell cycle regulators leading to impaired cell cycle progression. Although the role of Ubp14 in cell cycle regulation in yeast is poorly characterised, its human homolog, USP5 has been implicated in cell cycle control. Pan-cancer analyses of USP5 showed that elevated expression of USP5 correlates with tumour progression. To examine whether Ub^L8A^ mediated Ubp14 inhibition similarly affected the cell cycle progression in yeast, we performed an asynchronous FACS-based cell cycle analysis in the W303, ΔUbp14 and ΔUbp13 strains expressing either the wild-type or the mutant ubiquitin (Fig 3C). We observed an increased population of cells with 2C DNA content upon expression of Ub^L8A^ in W303 compared to cells with the wild-type ubiquitin suggesting a delayed G2/M phase progression. The ΔUbp14 strain alone or with Ub^WT^ or Ub^L8A^ overexpression, demonstrated a similar accumulation of cells in the G2/M phase due to the deletion of Ubp14 in these cells. This observation supported that Ub^L8A^ induced inhibition of Ubp14 in the yeast cells closely phenocopies the complete loss of Ubp14 function. Since we had observed slight accumulation of polyubiquitinated proteins in the ΔUbp13 strain expressing Ub^L8A^, we examined this strain for defects in progression of cell cycle. Similar to W303, the ΔUbp13 strain also demonstrated a decrease in the 1C/2C ratio (from ∼0.81 to ∼0.56) suggesting a delayed cell cycle progression due to an inhibition of the Ubp14 by Ub^L8A^ in this strain (data not shown). This suggested that Ubp13 is not the primary deubiquitinase targeted by Ub^L8A^.

To identify the phase of cell cycle progression affected by the mutant ubiquitin, we analysed the DNA in the synchronized W303 cells expressing Ub^WT^ or Ub^L8A^ over one generation of cell growth. Compared to the control cells, Ub^L8A^ expression in W303 showed a slower progression into S phase with lower 2C DNA over time (Fig 3C). We also observed a delayed exit from G2/M phase at ∼120 minutes in Ub^L8A^ expressing cells compared to wild-type ubiquitin which suggested that the Ub^L8A^ impaired the timely mitotic progression in the W303 cells. Together, these results provided a strong functional evidence that the Ub^L8A^ expression stabilized the Ubp14 and polyubiquitin chain complex that disrupts the ubiquitin homeostasis and compromises ubiquitin dependent cellular processes.

### L8A mutant ubiquitin exhibits enhanced interface interactions with USP5

While our biochemical assays showed a stronger association of Ub^L8A^ comprising polyubiquitin chains with the Ubp14 in the yeast cells, the structural basis of this interaction remained unclear. Hence, we performed an all-atom molecular dynamics (MD) simulations on the Ub: DUBs complex using the AMBER simulation suite (Case et al., 2025; Salomon et al., 2013) and AMBER ff19SB force fields (Tian et al., 2019). Since the crystal structure of Ubp14 was not available, simulations were initiated from the high-resolution structure of ubiquitin bound to USP5 (PDB: 3IHP). The system comprised of two USP5 molecules, each bound to a single ubiquitin (Fig 4A and 4B), enabling broad sampling of conformational space and a detailed characterisation of the complex.

**Figure 4:**
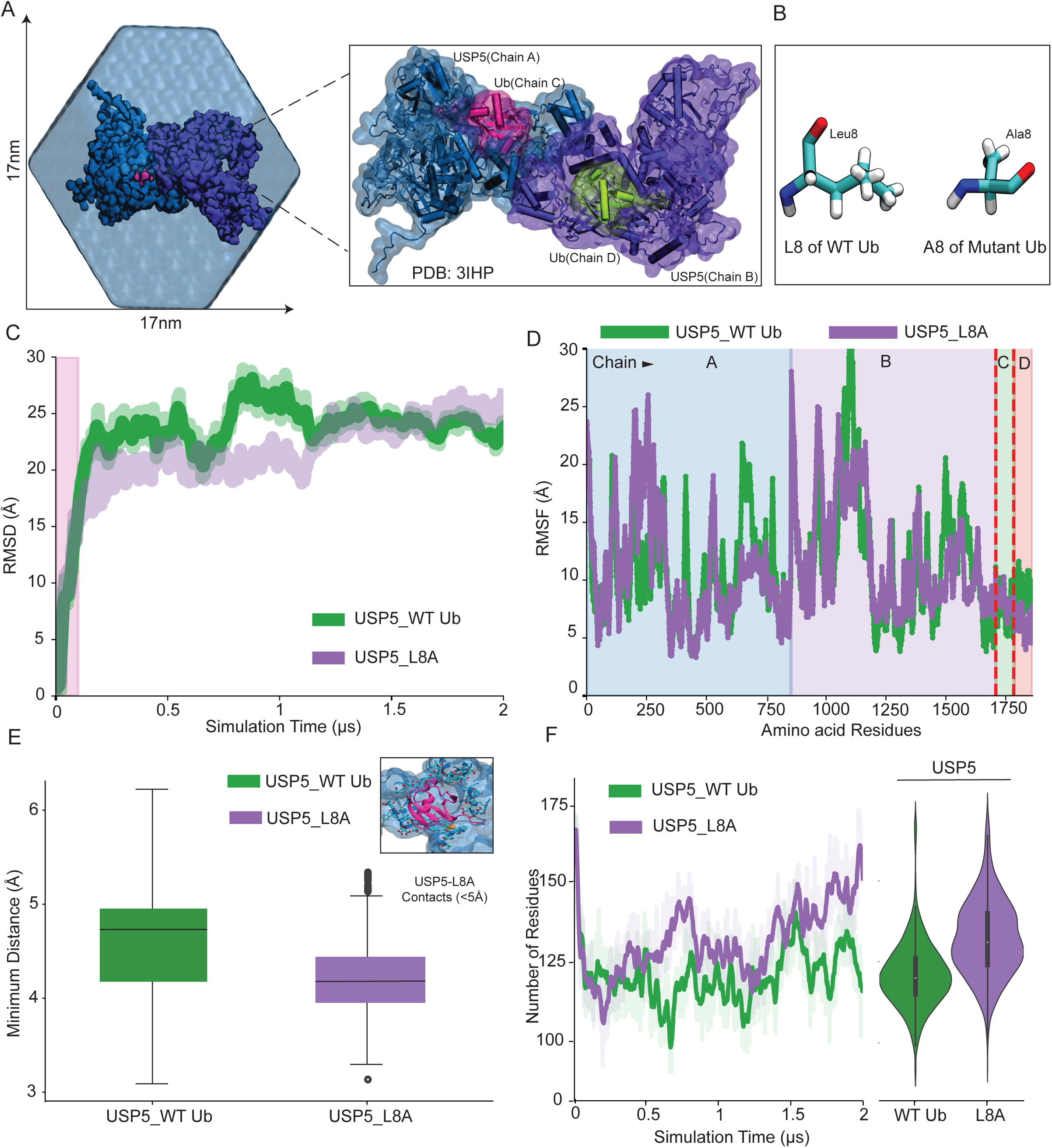
L8A mutant ubiquitin exhibits enhanced interface interactions with USP5. (A). Representative snapshot of the all-atom model of PDB ID: 3IHP (System 3) solvated in an explicit water box. A close-up view highlights the secondary structures of the two USP5 molecules (blue and violet) and two ubiquitin molecules (pink and green). (B). Chemical structures of leucine and alanine amino acids. (C). Root-mean-square deviation (RMSD) of the entire protein complex for Systems 3 and 4 as a function of simulation time. (D). Root-mean-square fluctuation (RMSF) plotted as a function of residue index. Residues 1–1710 correspond to USP5, whereas residues 1711–1862 correspond to ubiquitin. (E). Minimum distance between USP5 and ubiquitin over simulation time, averaged across two identical complexes. (F). Number of USP5 residues within 5 Å of ubiquitin, representing intermolecular contacts, averaged across two identical complexes.

We systematically analysed the molecular dynamics, structural stability and intermolecular interactions of the Ub^WT^ and Ub^L8A^, both in the unbound and as USP5 complex states. All-atom MD simulations was first performed for unbound ubiquitin and the mutant in explicit water (systems 1 and 2) and later in the complex with USP5 (systems 3 and 4; Fig 4A and 4B and Supplementary table 3). Each system was simulated for a total of 2μs which allowed us to evaluate long-timescale motions and subtle conformational rearrangements. Across all simulated system, the global protein fold remained stable throughout the trajectories (data not shown), however, we observed differences in the protein-protein interactions.

The molecular dynamics (MD) simulations for unbound forms revealed an increased structural flexibility and conformational heterogeneity in Ub^L8A^ relative to the Ub^WT^. This suggested that the leucine to alanine substitution altered the conformational ensemble of ubiquitin, potentially influencing its binding behaviour toward USP5. Analysis of the structural deviations in USP5:L8A complex revealed that Ub^L8A^ exhibited a lower overall root mean square deviation (RMSD) compared to the USP5:WT for majority of the simulation period (Fig 4C). This indicated an enhanced structural stability of the Ub^L8A^: USP5 protein complex during the initial phase of the simulation. Root mean square fluctuation (RMSF) analysis further revealed chain-specific effects of the bound wild-type or mutant ubiquitin in the USP5 dimer. Notably, L8A bound to the chain A of USP5 showed reduced fluctuations relative to the wild-type ubiquitin bound to the chain A, suggesting increased local stability in part of the complex. In contrast, L8A bound to the chain B of USP5 displayed slightly higher fluctuations than its wild-type counterpart (Fig 4D). These findings suggested that the L8A mutation imparts asymmetric effects on the two chains of the USP5 dimer, potentially altering inter-chain dynamics or local flexibility in a non-uniform manner.

To investigate the interaction dynamics between the wild-type ubiquitin or mutant and USP5, we measured the minimum distance between the two proteins in their bound state throughout the simulations. We observed that Ub^L8A^ maintained a significantly closer proximity to USP5 compared to the Ub^WT^: USP5 complex (Fig 4E). Given that single-residue substitutions can propagate altered binding effects to distal residues, we next assessed conformational changes in USP5 residues contacting the ubiquitin protein. The analysis revealed that the Ub^L8A^:USP5 complex had a greater number of residues within a close contact distance relative to the Ub^WT^: USP5 complex (Fig 4F), indicating an extensive interaction interface that likely contributes to the enhanced complex stability. Although the solvent accessible surface area (SASA) for bound and unbound forms of WT or mutant ubiquitin were comparable (data not shown). To identify representative conformational states, we performed hierarchical agglomerative clustering of MD trajectories for the Ub^WT^ and Ub^L8A^ complexes (systems 3 and 4). Ten dominant clusters were identified for each system, and the most populated cluster from each trajectory was selected for a detailed interaction analysis. Structural assessment of these representative conformations using PDBSUM (Laskowski et al., 2018) revealed that the Ub^L8A^:USP5 complex formed a substantially higher number of stabilising interactions, including hydrogen bonds, salt bridges, and non-bonded contacts, relative to the WT complex (data not shown). Further the L8A mutation induces a compactness when bound to the USP5 opposed to its unbound state, restricting the conformational dynamics required for appropriate ubiquitin chain processing by USP5. Taken together, these results revealed that the Ub^L8A^:USP5 complex exhibited an increased number of non-covalent interactions, demonstrated a more compact conformation, and showed reduced structural deviations, collectively indicating enhanced stability relative to the Ub^WT^:USP5 complex.

### L8A mutant ubiquitin intrinsically inhibits deubiquitinase activity of Ubp14

The in vivo functional analysis of Ub^L8A^ expression clearly demonstrated that the mutant comprising polyubiquitin chains associated strongly with the Ubp14 to inhibit its deubiquitinase activity but whether the unconjugated mutant can also inhibit the deubiquitinase or the mutant’s inhibitory effect is strictly chain dependent remained unclear. To address this, we performed the in vitro deubiquitinase assay with the purified proteins. We purified the wild-type ubiquitin, the L8A mutant ubiquitin and the Ubp14 and assessed the structural integrity of the mutant ubiquitin protein prior to the activity assay. We observed that Ub^L8A^ protein retained its secondary structure and thermal stability similar to the Ub^WT^ suggesting that the leucine to alanine substitution did not perturb the global folding of the protein also shown by the ΔG analysis of this mutant (Supplemental table 2).

Ub-AMC cleavage assay revealed that Ub^L8A^ inhibited both the Ubp14 and USP5 in a dose-dependent manner, with an estimated IC_50_ of ∼1.44 µM. In contrast, Ub^WT^ little inhibition even at the high substrate concentration with an IC_50_ of >3 µM for both Ubp14 and Usp5 (Fig 5A and 5B). This suggested that Ub^L8A^ had a markedly stronger association with both the DUBs. We also observed minimal fluorescence signals of substrate processing by Ubp14 when we used the Ub^L8A^ with terminal diglycine motif deleted in this mutant sequence which indicated that Ub^L8AΔGG^ also strongly associates with Ubp14 similar to Ub^L8A^ (Fig 5C). Hence, the inhibitory effect of Ub^L8A^ is mediated primarily by its hydrophobic patch and not by the canonical C-terminal glycine for binding in the in vitro assay where other components that mediate ubiquitin conjugation are also not included (Fig 5C). This result suggests that Ub^L8A^ does not impair substrate recognition by Ubp14 rather it interferes with the productive catalysis of linked ubiquitin by occupying the catalytic pocket of Ubp14.

**Figure 5:**
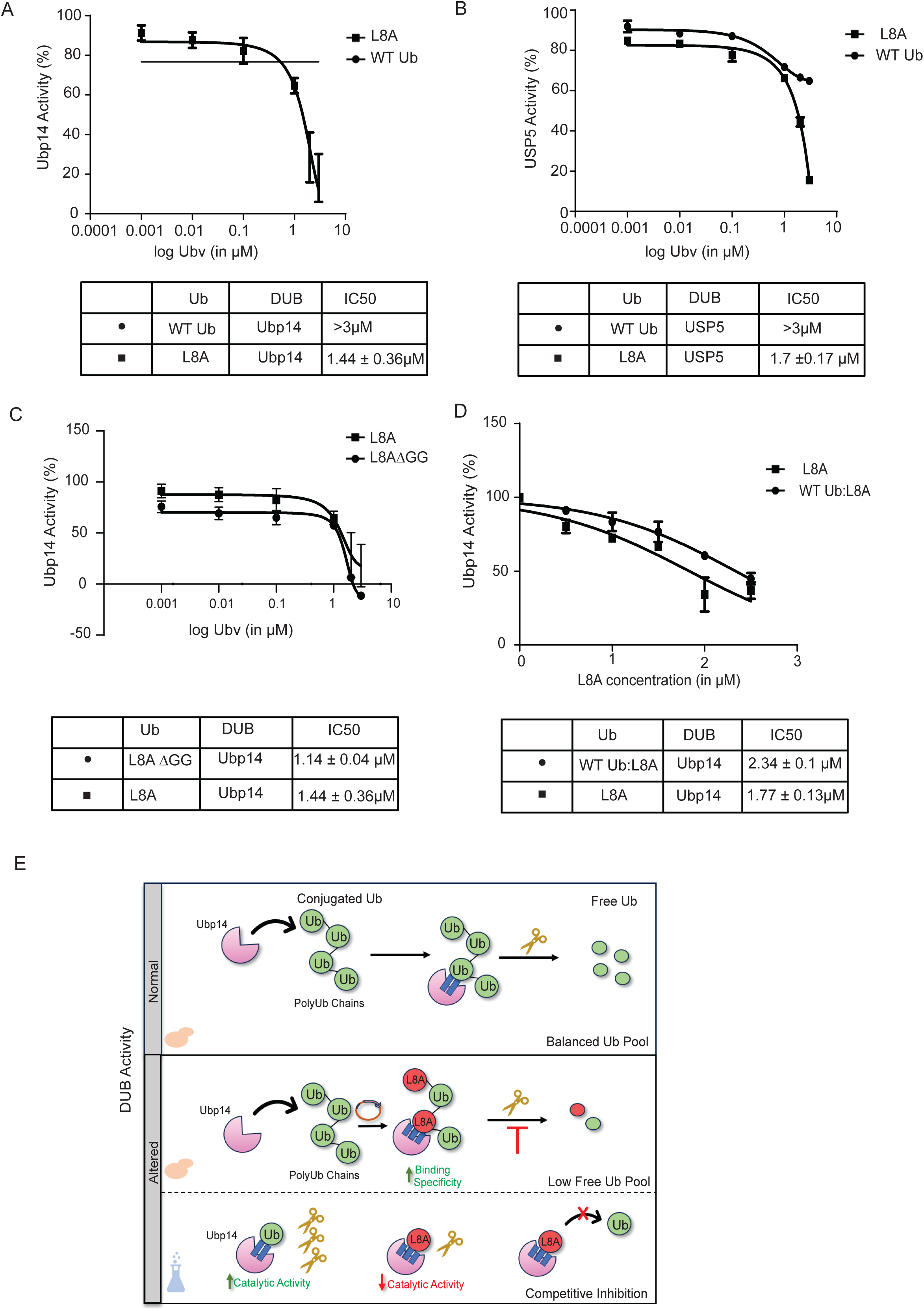
L8A mutant ubiquitin intrinsically inhibits deubiquitinase activity of Ubp14. (A). Dose–response curves for inhibition of Ub-AMC hydrolysis by Ubp14 in the presence of ubiquitin mutant L8A (squares) or wild type (circles). Normalized activity was determined relative to activity in the absence of inhibitor. Data are presented as mean ± SEM (n = 3). The IC50 value was calculated as the concentration of ubiquitin mutant that reduced proteolytic activity by 50%. (B). Dose–response curves for inhibition of Ub-AMC hydrolysis by USP5 in the presence of ubiquitin mutant L8A (squares) or wild type (circles). Normalized activity was determined relative to activity in the absence of inhibitor. Data are presented as mean ± SEM (n = 3). The IC50 value was calculated as the concentration of ubiquitin mutant that reduced proteolytic activity by 50%. (C). Dose–response curves for inhibition of Ub-AMC hydrolysis for Ubp14 in the presence of ubiquitin mutant L8A (squares) or L8AΔGG (circles). Normalized activity was determined relative to activity in the absence of inhibitor. Data are presented as mean ± SEM (n = 3). The IC50 value was calculated as the concentration of ubiquitin mutant that reduced proteolytic activity by 50%. (D). Dose–response curves for inhibition of Ub-AMC hydrolysis for Ubp14 in the presence of ubiquitin mutant L8A (squares) or L8A in presence of WT Ub (circles). Normalized activity was determined relative to activity in the absence of inhibitor. Data are presented as mean ± SEM (n = 3). The IC50 value was calculated as the concentration of ubiquitin mutant that reduced proteolytic activity by 50%. (E). Model illustrating altered DUB activity upon expression of the L8A ubiquitin mutant. Under normal conditions, Ubp14 binds unanchored polyubiquitin chains (green circles) and cleaves them to maintain the cellular free ubiquitin pool. When the L8A mutant (red circles) is expressed, it binds strongly to Ubp14 (blue lines), enters the ubiquitin pool, and forms mixed chains with WT Ub, thereby disrupting DUB activity and reducing free ubiquitin levels. In vitro, a single L8A molecule can strongly bind Ubp14 and interfere with its catalytic activity.

To mimic our in vivo experimental conditions where the inhibitory effects of Ub^L8A^ are exerted in the presence of the endogenously expressed ubiquitin, we assessed L8A mediated inhibition in presence of the wild-type ubiquitin. Interestingly, Ub^L8A^ retained its inhibitory activity even under these competitive conditions, with an IC_50_ of ∼1.7 µM, demonstrating that the mutant ubiquitin can dominantly and competitively occupy the DUB binding site (Fig 5D). Collectively, these results establish that the engineered L8A variant stabilises the DUB–ubiquitin complex and functions as a potent, selective inhibitor, providing a mechanistic explanation for its dominant effects observed in vivo.

Our findings demonstrate (Fig 5E) that under physiological conditions, Ubp14 deubiqutinase efficiently cleaves ubiquitin molecules from the free or substrate bound polyubiquitin chains, maintaining a homeostatic pool of free ubiquitin. However, when the L8A mutant ubiquitin, as a part of the polyubiqutin chain in vivo or as free ubiquitin in vitro, preferentially engages with the Ubp14, it interferes with the normal the ubiquitin processing function of the Ubp14. This competitive inhibition reduces free ubiquitin as shown both in vitro and in vivo, highlighting that L8A acts as a tunable modulator of ubiquitin pools in the cells.

## Discussion

This study establishes that leucine 8, a hydrophobic patch residue of ubiquitin, is a critical determinant of Ubp14 deubiquitinase function in cells. We demonstrate that under overexpression conditions, a specific substitution of leucine 8 to alanine in ubiquitin converts ubiquitin into a Ubp14 binding-competent yet catalytically inert substrate thereby behaving as a dominant negative inhibitor of Ubp14. Mutant ubiquitin is efficiently incorporated into the polyubiquitin chains in the cells that associates with enhanced intermolecular interactions with Ubp14 to prevent the catalytic processing of the polyubiquitin chain within the active site of Ubp14. Interestingly, these findings reveal that even subtle perturbations within the hydrophobic patch of ubiquitin exert disproportionately large effects on DUB specificity, uncovering a previously underappreciated layer of residue-specific regulation within the ubiquitin interaction interface.

Despite ubiquitin having minimal structural specialization for its interaction with a wide range of cellular proteins, we were able to identify a residue-specific substitution that enables specific functional regulation of ubiquitin. No other substitution at the L8 position showed similar inhibition of Ubp14 function in the cells under our experimental conditions. The L8A mutation did not alter the stability of ubiquitin but modified the Ubp14 and ubiquitin interface to promote stable but non-productive engagement of Ubp14 with the mutant ubiquitin. This strong association of Ubp14 by Ub^L8A^ trapped Ubp14 quenching it from the cellular pool for cleaving the wild-type like polyubiquitin chains. This “trapping effect” of Ubp14 by the Ub^L8A^ showed a dosage-dependent phenotype where overexpression of Ubp14 relieved the toxic effects of the mutant in the cells. Interestingly, our in vitro results showed that the unconjugated, free mutant ubiquitin (Ub^L8AΔGG^) also inhibited the Ubp14 activity (Fig 5C), although these effects are likely masked in the complex cellular environment where the conjugated ubiquitin forms are a prominent specie. We demonstrated that the dominant negative effects of L8A have measurable physiological consequences revealing that cellular homeostasis is highly sensitive to a balanced pool of free and conjugated ubiquitin. Conclusively, our study revealed a finely tuneable node of ubiquitin function which is obscure with traditional approaches that prioritize strongly toxic or loss of function mutations. This study strengthened our earlier findings that single residue perturbations can yield dominant negative effects underscoring the functional sensitivity of ubiquitin interface (Padhy et al., 2023)

DUBs have distinct and complex mechanisms of recognizing substrate conjugated ubiquitin molecules (Lange et al., 2022). The selective inhibition of Ubp14 by L8A mutant can be rationalized by considering that as opposed to most DUBs, Ubp14 comprises of four ubiquitin binding sites to recognize the sequentially associated ubiquitin molecules in distinct chain topologies (Gao et al., 2024). While most DUBs utilize the C-terminal end and the hydrophobic patch of the substrate-bound ubiquitin to associate with it while positioning the cleavable isopeptide bond between ubiquitin and substrate for catalytic cleavage, Ubp14 uniquely follows this interaction mode with multiple ubiquitin units in a consecutive manner. This extended recognition mechanism requires a precise alignment of adjacent ubiquitin units which imposes a strong dependence of Ubp14 on both chain topology as well as hydrophobic patch geometry, making Ubp14 sensitive to subtle perturbations in ubiquitin structure. Previous studies have shown that single residue perturbations in DUBs or in the ubiquitin protein modulate the substrate or linkage specificities by altering the binding affinity of molecules in the complex (Morrow et al., 2018; Phillips et al., 2013; Reyes-Turcu et al., 2009b; Ronau et al., 2016). In contrast, L8A mutant ubiquitin described in this study enhanced binding with Ubp14 while impairing its catalytic turnover, thereby uncoupling ubiquitin recognition from ubiquitin processing functions of Ubp14.

The hydrophobic patch of ubiquitin comprising of Leu8, Ile44 and Val70 residues has been implicated in mediating interactions with numerous cellular proteins (Dikic et al., 2009a; Haririnia et al., 2008; Harper and Schulman, 2006; Singh et al., 2017). Structural and biochemical analyses demonstrate that the majority of the ubiquitin receptors interact via Ile44 (Dikic et al., 2009b; Husnjak and Dikic, 2012). It is this primary interacting residue of this patch with solvent oriented side chain while Leu8 side chain is directed to the interior of this patch but interestingly, leucine appears to be a more buried residue in the interface of ubiquitin with DUBs in majority of the co-crystal structures available (Fig. 1C). Alanine mutagenesis studies indicated that all the three residues of the patch contribute to ubiquitin-partner protein interaction, however, their effects are not equivalent, with I44A having more pronounced deleterious phenotypes compared to Leu8 or Val70 (Kang et al., 2003; Singh et al., 2017; Sloper-Mould et al., 2001b). We demonstrated that unlike Ile44, the L8A mutation did not abolish its partner binding but instead it reduced the efficiency of ubiquitin dependent processes. This distinction is more evident under the over-expression conditions used in our experiments and led to selective impairment of the Ubp14 functions. This study supported the role of Leu8 as a peripheral residue of the patch for precise spatial organisation of ubiquitin for efficient catalysis rather than serving as a primary anchoring point.

The selective inhibition of Ubp14 and its human homolog, the USP5 deubiquitinase, has important cellular implications particularly in tumor progression and cancer cell survival where USP5 is known to be upregulated (Jiang et al., 2025; Kaistha et al., 2017; Yan et al., 2023). Small molecule inhibitors of the catalytic domain of USP5 have been developed but owing to the conservation of the targeting domain across the USP family, these inhibitors have limited selectivity for USP5. In contrast, the inhibition of USP5 by the L8A ubiquitin mutant is functionally distinct, where L8A containing polyubiquitin chain binds strongly with the USP5 ubiquitin binding domains however these higher order interactions are inefficient for cleaving the polyubiquitin chain. As this mode of inhibition is strongly dependent on ubiquitin chain architecture, and binding of multiple ubiquitin units to USP5, the L8A mutant offers higher potential for selectivity compared to conventional inhibitors. Engineered high affinity binders to DUBs have been generated through direct competition of ubiquitin variants for enzyme binding (Guo et al., 2021; Tang et al., 2023, 2021; Veggiani et al., 2022). Notably, this approach has identified the embedded plasticity within the ubiquitin interaction surface. These ubiquitin variants incorporate multiple substitutions to enable strong binding towards specific DUBs, however, their optimised affinities might alter in the complex cellular environment. We explore how minimal perturbations of the ubiquitin native structure influence both the ubiquitin-ubiquitin and ubiquitin-enzyme interactions in a physiological context. This study offers a conceptual framework for developing selective DUBs inhibitors by disrupting substrate geometry rather than catalytic activity only.

While in this study we present the mechanistic insights into how L8A serves as a substrate inhibitor for the cellular functions of Ubp14, several aspects remain unexplored. Molecular simulation analysis demonstrated the local interaction changes induced by the mutation, however, this analysis is limited to the positioning of a single ubiquitin molecule in USP5. It will be interesting to explore the altered chain topology of a polyubiquitin chain having L8A mutant ubiquitin. High-resolution structural information will be required to understand the changes in positioning of L8A comprising polyubiquitin chain within the four ubiquitin binding domains of Ubp14. Biochemical and proteomics analysis under the mutant overexpression conditions can help to determine if specific chain linkages are preferentially enriched under these conditions. USP5 is particularly sensitive to ubiquitin geometry as it uniquely engages with multiple ubiquitin moieties in a polyubiquitin chain. We observed L8A induced inhibition of USP5 functions in complementary yeast cellular assay and in vitro assay, however, this selectivity can be confirmed in mammalian cells which display a broader landscape of deubiquitinases.

Conclusively, we demonstrated a cryptic node of proteostasis regulation residing in the hydrophobic patch of ubiquitin that enables it to fine-tune the enzymatic activity of Usp14 deubiquitinase. Dominant negative mutants of ubiquitin offer a unique approach to dissect the complex ubiquitin signalling pathway as shown for the L8A mutant that uncouples substrate binding efficiency from enzyme catalysis.

## Materials and Methods

### Yeast strains and plasmid constructs

Most strains used in this study were derived from the *Saccharomyces cerevisiae* W303 MATa background. Individual DUB deletion strains were generated by homologous recombination using standard gene deletion protocols (Rothstein, 1991). Ubiquitin variants used for screening were produced by site-directed mutagenesis and cloned under the GAL1 promoter into the pRS424 or pRS413 vectors (Addgene, 40235). The *Ubp14* gene was amplified from W303 genomic DNA and subcloned into the URA3-marked pRS316-Cu-GFP backbone (Addgene,1037). Human USP5 was cloned into the same pRS316 vector using cDNA isolated from V6R prostate cancer cells as the template. Primer sequences are listed in Table 1.

**Table 1.**
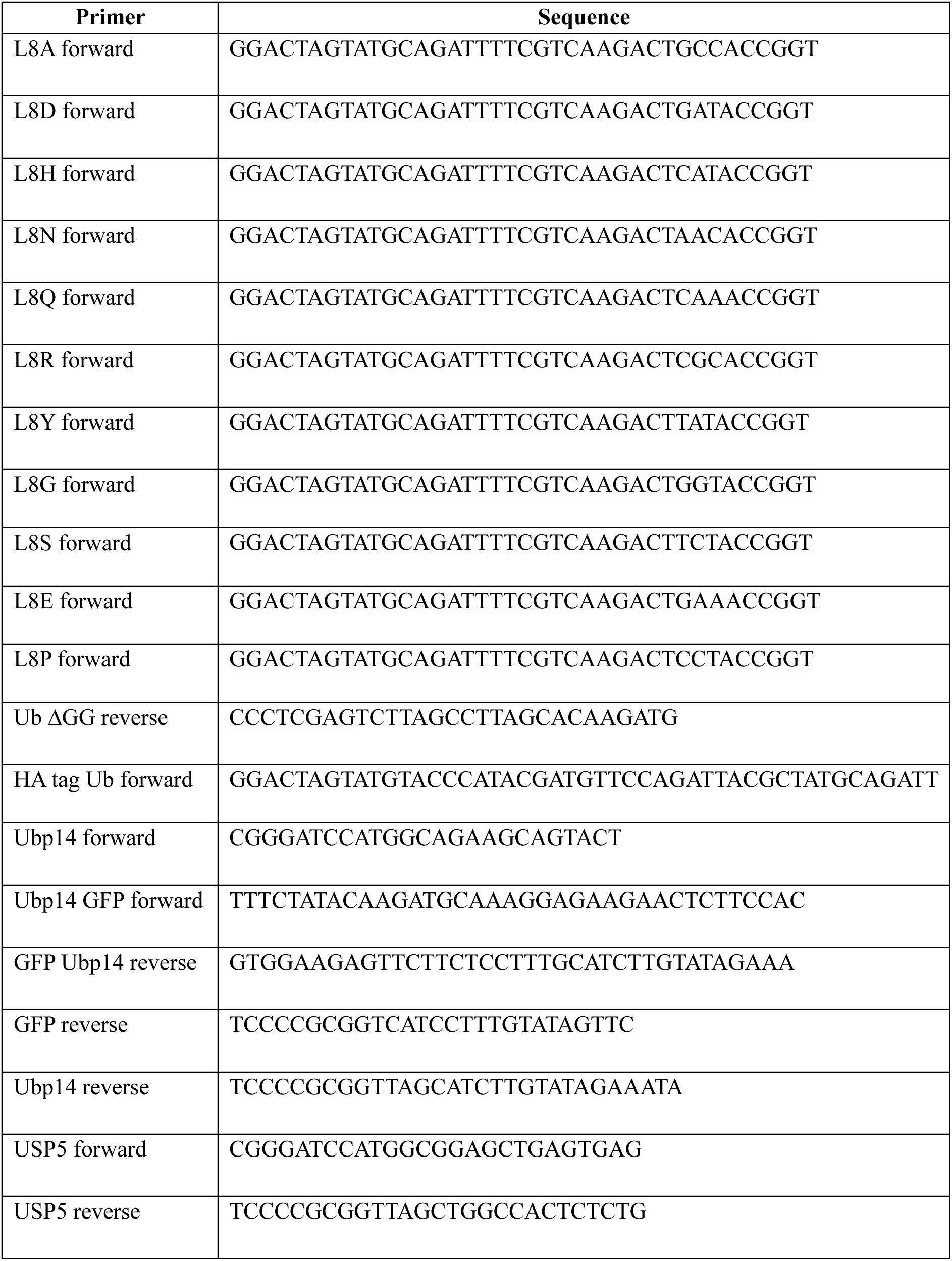
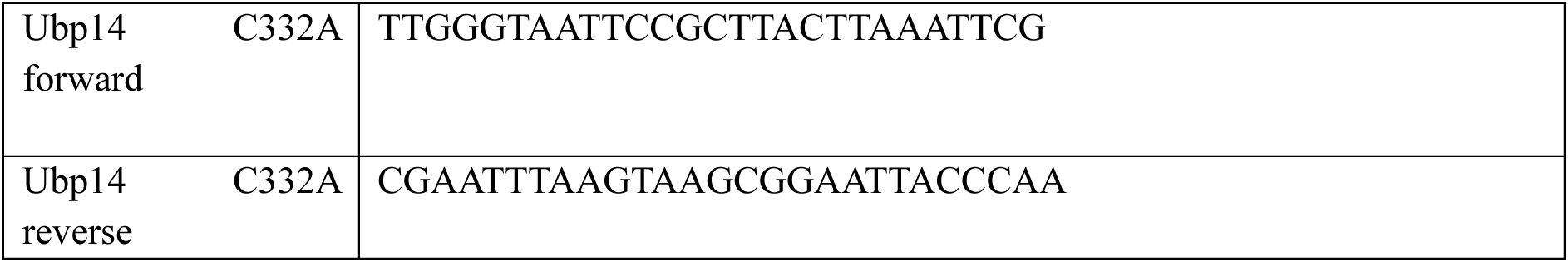
Primers used in this study.

For ubiquitin shutoff experiments, we used the Sub328 strain, which expresses a single copy of ubiquitin under the galactose-inducible promoter and can be repressed upon shifting to dextrose. Ubiquitin variants were cloned into the p427-GPD vector and transformed into the shutoff strain using the lithium acetate method as previously described (Roscoe et al., 2013b). All yeast strains and plasmids used in this study are listed in Table 2 and Table 3, respectively.

**Table 2.** Yeast strains used in this study.

| Strains | Genotype | Source |
| --- | --- | --- |
| S.cerevisiae W303 Mat-a | leu2-3,112 trp1-1 can1-100<br>ura3-1 ade2-1 his3-11,15 | This study |
| Sub328 shut-off | ubil-δ1::trp1 ubi2-δ2::ura3<br>ubi3δub-2 ubi4-δ2::leu2 +<br>[pub146] + [pub100]) | This study |

**Table 3.** Plasmid constructs used in this study.

| Recombinant DNA | Source |
| --- | --- |
| pRS424 Gal WT Ub | This study |
| pRS424 Gal L8A Ub | This study |
| pRS424 Gal L8D Ub | This study |
| pRS424 Gal L8H Ub | This study |
| pRS424 Gal L8M Ub | This study |
| pRS424 Gal L8N Ub | This study |
| pRS424 Gal L8R Ub | This study |
| pRS424 Gal L8Q Ub | This study |
| pRS424 Gal L8Y Ub | This study |
| pRS424 Gal L8G Ub | This study |
| pRS424 Gal L8S Ub | This study |
| pRS424 Gal L8E Ub | This study |
| pRS424 Gal L8P Ub | This study |
| pRS424 Gal HA WT Ub | This study |
| pRS424 Gal HA L8A Ub | This study |
| pRS424 Gal HA L8N Ub | This study |
| pRS424 Gal HA L8Q Ub | This study |
| pRS424 Gal HA L8H Ub | This study |
| pRS413 Gal GFP | Addgene (40235) |
| pRS413 Gal WT Ub | This study |
| pRS413 Gal L8A | This study |
| pRS413 Gal L8S | This study |
| pRS316 Cu GFP | Addgene (1037) |
| pRS316 Cu GFP Ubp14 | This study |
| pRS316 Cu USP5 | This study |
| pET21 His6X WT Ub | Bolon Lab |
| pET21 His6X L8A Ub | This study |
| pET21 His6X L8A ΔGG Ub | This study |
| pET21 His6X Ubp14 | This study |
| pET21 His6X Ubp14 C332A | This study |
| <u>pUB23-P-beta-gal</u> | Addgene (8817) |
| <u>pUB23-L-beta-gal</u> | Addgene (8820) |
| p427 GPD WT Ub | This study |
| p427 GPD L8A Ub | This study |

### Yeast growth rate and mutant expression analysis

To estimate the growth rate of selected L8 ubiquitin mutants in yeast, mutant plasmids were transformed and expressed in yeast as described previously (Padhy et al., 2023). Briefly, transformants were selected on SD-Trp plates and grown in SD-Trp to saturation. Cultures were re-inoculated, grown to early log phase, washed, and shifted to SRafGal-Trp medium (1% galactose) for 6 h induction. OD-normalised cultures were serially diluted and spotted onto SD-Trp and SRafGal-Trp plates, which were incubated at 30 °C for 2-3 days. All plate images were analysed with ImageJ software.

Ubiquitin point mutants were generated in the p427GPD vector and transformed into the Sub328 shutoff strain as previously described. Yeast transformants were grown in synthetic complete (SC) medium containing G418 and supplemented with 1% raffinose and 1% galactose (RGal). For plate-based growth assays, cells were pre-grown for 6 h in SC + G418 containing 2% dextrose, serially diluted, and spotted onto SC + G418 plates containing either RGal or dextrose. Plates were incubated for 2 days at 30 °C.

For liquid growth assays, strains were grown to mid-log phase in SC + G418 RGal medium, shifted to SC + G418 dextrose medium, and cultured for 16 h at 30 °C to shut off and deplete endogenous wild-type ubiquitin. Cultures were then diluted to early log phase (OD₆₀₀ = 0.1) in fresh SC + G418 dextrose medium and grown at 30 °C, with OD₆₀₀ monitored over time to assess growth kinetics. Ubiquitin levels were analysed by western blotting using an anti-ubiquitin antibody and chemiluminescent detection.

The HA-tagged ubiquitin constructs were cloned into pRS424 Gal yeast expression plasmid and were subsequently transformed into W303 cells, which retain endogenous ubiquitin expression. Yeast cell lysates were prepared as previously described. Ectopic expression of HA tagged ubiquitin was analyzed by western blotting using an anti-HA antibody (Antibodies.com; HA.C5, A85278).

### Yeast phenotypic analysis of ubiquitin mutant overexpression in DUB deletion strains

For screening of selected mutants in different DUB deletion strains, wild-type and mutant ubiquitin plasmids were transformed into different DUB deletion strains using similar methods. Serial dilution assay on SD-his and SRafGal-His media was performed for each strain containing the ubiquitin mutants after 6h of Gal induction. Growth of each strain expressing the mutants was measured as a fold change to wild-type expression in the same genetic background. For DUB overexpression studies, pRS316-Cu-Ubp14 and pRS316-Cu-USP5 were transformed into W303 and ΔUbp14 using similar methods. Individual colonies were inoculated and grown in SD-Ura media to prepare competent cells using 0.1M lithium acetate. Selected ubiquitin mutants were transformed into these cells and plated onto SD-His-Ura media the next day for the selection of cells containing both the DUB and the ubiquitin mutant. To check the mutant effect on overexpressed dubs, copper sulfate (50 μM) was added to the glucose medium at log phase and grown for 6 hours. The cells were then pelleted and washed with autoclaved double-distilled water to remove trace amounts of glucose and resuspended in synthetic media (SRafGal-His-Ura) having 1% galactose as the sugar source to induce the expression of mutants. After 3 hours of galactose induction, 10-fold serial dilutions of the samples were plated on SD-His-Ura and SRafGal-His-Ura+Cu plates and were grown for 2-3 days at 30 ^ο^C to monitor colony growth.

### Western blotting

Yeast sample preparation, lysis, and western blotting were performed as previously described (Padhy et al., 2023). Briefly, equal amounts of lysate were resolved by SDS–PAGE, transferred to PVDF membranes, blocked in 4% BSA, and probed with the appropriate antibodies. Polyubiquitin levels were detected using anti-ubiquitin (Santa Cruz-P4D1) at 1:1000 dilution. DUB and substrate expression were assessed using antibodies against GFP (Sigma Aldrich, 11814460001) and USP5 (Santa Cruz, C-11) at 1:2000, followed by HRP-conjugated secondary antibodies and chemiluminescent detection. Band intensities were quantified in ImageJ (NIH) and normalized to the corresponding loading control.

### Flow cytometry

Yeast cells after induction were harvested and fixed in 70% ethanol and stained with 16ug/ml of propidium iodide as previously described (Rosebrock, 2017). Data were collected on a Becton Dickinson LSRFortessa cell analyzer. Data were analyzed using FlowJo software (FlowJo, LLC). In arrest-release experiments, cells were arrested at the G1 phase using α-factor (Sigma Aldrich, T6901) at a final concentration of 240nM and then released after 3 hours by centrifugation and growing the cells in 1% galactose selection media. Samples were then collected at indicated time points and analyzed similarly.

### Cycloheximide chase assay

W303 cells harbouring the wild-type ubiquitin and L8A individually were transformed with plasmids encoding Ub-P-beta-gal and Ub-L-beta-gal (Addgene, 8817; 8820). Single transformants were grown in double dropout synthetic liquid media SD-Trp-Ura to saturation. From this saturated culture, 1% of the cells were diluted in the same non-inductive media, grown to the early log phase, pelleted, and washed with SRafGal-Trp-Ura to remove any amount of dextrose. The pellet was resuspended in induction media SRafGal-Trp-Ura and grown to mid-log phase, and then protein synthesis was inhibited by adding 50 ug/ml cycloheximide (Sigma Aldrich, C7698). Samples were collected at 0, 5, 10, and 20 min and analysed by western blotting. The lysates were probed with an anti-β-galactosidase primary antibody (Invitrogen, A11132)) at a 1:2000 dilution in 1X TBST.

### Co-Immunoprecipitation assay

Yeast cells expressing Ubp14-GFP and either wild-type or L8A ubiquitin were grown in selective media to mid-log phase and induced with 1% galactose. Cells were lysed in IP buffer (50 mM Tris-HCl pH 7.5, 150 mM NaCl, 1% Triton X-100, protease inhibitors) using glass bead lysis. Clarified lysates were incubated with Sepharose beads for 2 hours at 4°C. Beads were washed extensively, and bound proteins were eluted with SDS sample buffer. Eluates were analyzed via SDS-PAGE followed by immunoblotting with anti-ubiquitin and anti-GFP antibodies to detect co-immunoprecipitated ubiquitin.

### Protein Purification

Expression and purification of His₆-tagged wild-type and mutant ubiquitin from the pET21 vector were performed as previously described (Padhy et al., 2023). Briefly, constructs were expressed in BLR (DE3) *E. coli* under ampicillin selection, purified using Cobalt–NTA resin, and dialyzed into potassium phosphate buffer. Following elution, 10 mM DTT was added and samples were incubated for 1 h at 4 °C to ensure reduction of disulfide bonds.

Wild-type or mutant Ubp14 was also expressed similarly in the pET21 bacterial expression vector with protein expression induced with 0.5mM IPTG for 18h at 18^ο^C. Cells were pelleted and harvested in lysis buffer A (50 mM sodium phosphate buffer pH 7.2, 300 mM Nacl, 0.01% Triton X, 5% glycerol, 1mM beta-mercaptoethanol and 10 mM imidazole). The insoluble fraction containing the protein was resuspended in denaturing lysis buffer A with 8M urea and lysed by bead-beating at 4 °C. The lysate was clarified by centrifugation (12,000 × g, 10 min, 4 °C), and the supernatant containing denatured Ubp14 was transferred to a clean tube. To refold the protein, the denatured supernatant was subjected to stepwise urea reduction by sequential dialysis with lysis buffer A to final urea concentrations of 6 M, 4 M, 2 M, and 0 M, allowing gentle mixing for 3 hours at each step. The fully refolded lysate was then clarified again (12,000 × g, 10 min, 4 °C) to remove aggregates and applied to Co-NTA agarose pre-equilibrated in native binding buffer (50 mM sodium phosphate buffer pH 7.2, 300 mM NaCl, 0.01% Triton X, 5% glycerol and 10 mM imidazole). After binding for 1 h at 4 °C with gentle rotation, the resin was washed with 10 column volumes of binding buffer, followed by washing with high-stringency buffer (50 mM sodium phosphate buffer pH 7.2, 300 mM NaCl, 0.01% Triton X, 5% glycerol and 20 mM imidazole). Bound Ubp14 was eluted with 250 mM imidazole 50 mM sodium phosphate buffer pH 7.2, 150 mM Nacl, 0.01% Triton X and 5% glycerol, concentrated and buffer-exchanged into storage buffer (50mM sodium phosphate, 150mM NaCl). Purity was assessed by SDS–PAGE. Estimation of protein concentrations was carried out by calculating the absorption at 280 nm (Nanodrop 1000, Thermo Scientific, Rockford, IL, USA. Purified human USP5 protein was purchased from Sino Biological (12772-H08B).

### Circular dichroism

Circular dichroism (CD) spectra were acquired on a Jasco J-810 spectropolarimeter using a 1-mm pathlength quartz cuvette, with a 0.2-nm step size, 1-nm bandwidth, and a scan speed of 50 nm/min. Each spectrum represents the average of three accumulations and was baseline-corrected using the corresponding buffer spectrum. CD spectra of 5 μM wild-type or mutant ubiquitin, or Ubp14, were recorded in assay buffer (50 mM sodium phosphate, pH 7.5) over a wavelength range of 190–260 nm at 25 °C. Thermal unfolding profiles were obtained by monitoring the ellipticity at 220 nm while increasing the temperature from 20 to 90 °C at a rate of 1 °C/min using a Peltier temperature controller (Jasco PTC-423S). Ellipticity values were plotted against temperature and smoothed using a Loess fit in Prism 9.

### In vitro deubiquitinase assay

DUB enzymatic activity and inhibition assays were performed at room temperature using the fluorogenic substrates Ub-AMC (Boston Biochem, U-550). Reactions were carried out in assay buffer (50 mM PBS, pH 7.4, 0.01% Tween-20, 10 mM DTT) containing either 200nM of Ubp14 or USP5 and serial dilutions of Ub-AMC. AMC fluorescence (excitation 345 nm, emission 445 nm) was monitored for 30–60 min using the Variskon Lux plate reader (Thermofisher scientific). Initial reaction velocities were determined in duplicate from the linear increase in AMC fluorescence over time and converted to molar units using an AMC standard curve. Velocity versus substrate concentration data were fitted by nonlinear regression in GraphPad Prism 9 using the Michaelis–Menten equation to determine *K_M_*and *V*_max_. The catalytic constant (*k*_cat_) was calculated as *k*_cat_ = *V*_max_/*E*]_0_.

For inhibition assays, reactions were performed in assay buffer containing Ub-AMC (1 µM) as substrate. Serial dilutions of ubiquitin variants were preincubated with Ubp14 or USP5 for 30 min at room temperature. Proteolytic activity was monitored for 30 min in duplicate reactions. IC₅₀ values were determined by nonlinear regression in GraphPad Prism 9 from plots of UbV concentration versus percentage inhibition of Ubp14 and USP5 activity.

### Structural data analysis

Structural analyses and molecular images were generated using PyMOL (Schrödinger). The list of ubiquitin co-crystal structures with UPS components used for analysis was obtained from the RCSB PDB database and was manually curated to exclude structures of identical protein complexes (keeping only the highest resolution structure in each case) (Table S1). The average surface area buried at structurally characterised ubiquitin interfaces for the I44 hydrophobic patch residues was calculated using PyMOL. The crystal structure of ubiquitin (PDB: 1UBQ) was used for the mutational scanning analysis as position L8 of ubiquitin. Initially, the structure was processed using the RepairPDB function in FoldX to optimize side-chain conformations and minimize structural energy. Next, all possible amino acid substitutions at position L8 were generated using the PositionScan function in FoldX. The change in folding free energy (ΔΔG) for each mutant relative to the wild-type protein was calculated to assess the impact of substitutions on ubiquitin stability (Table S2).

USP5-Ubp14 sequence alignment was also performed using PyMOL. The Ubp14-Ub co-crystal structure was generated using AlphaFold 3 and aligned with the co-crystal structure of USP5-Ub (PDB ID: 3IHP) to assess the ubiquitin binding region.

### Molecular dynamics simulation analysis

All-atom molecular dynamics (MD) simulations were performed using the AMBER22 simulation suite (Case et al., 2022) with periodic boundary conditions. Long-range electrostatics were treated using the particle mesh Ewald (PME) method (Darden et al., 1993). Simulations were run with the CUDA-enabled pmemd.cuda module (Salomon-Ferrer et al., 2013) on an RTX4080 GPU workstation (Yamuna). The systems were maintained at constant pressure and temperature using a Monte Carlo barostat with isotropic position scaling (Berendsen et al., 1984) and a Langevin thermostat with a collision frequency of 1 ps⁻¹ (Loncharich et al., 1992). Joung and Cheatham ion parameters (Joung and Cheatham, 2008) with the TIP3P water model (Jorgensen et al., 1983) were used to describe interactions among Mg²⁺, Na⁺, Cl⁻, and water molecules. A cutoff of 8 Å was applied for van der Waals and short-range electrostatic interactions. All simulations employed a 2fs integration time step. The SETTLE algorithm (Miyamoto and Kollman, 1992) was applied to constrain water molecules, while the RATTLE algorithm (Andersen, 1983) constrained covalent bonds involving hydrogen atoms. System coordinates were saved every 20 ps for analysis. Trajectory analysis and post-processing were performed using VMD (Humphrey et al., 1996) and CPPTRAJ (Roe and Cheatham, 2013).

Simulations were initiated from the experimentally resolved crystal structure of the covalently linked ubiquitin–USP5 complex (PDB ID: 3IHP). The full complex contains 1,860 residues; however, only 1,501 residues were resolved in the crystal structure. The remaining 359 residues were modelled using SWISS-MODEL (Waterhouse et al., 2018) and fitted into the crystal structure using an in-house TCL script. All-atom topologies and parameters were generated using the xleap module of AMBER22, with the ff19SB force field (Tian et al., 2020) describing bonded and non-bonded interactions. Equilibrium MD simulations were carried out for four systems: (1) wild-type ubiquitin in water and ions, (2) L8A mutant ubiquitin in solution, (3) wild-type ubiquitin bound to USP5, and (4) L8A ubiquitin bound to USP5 (Table S3). Systems were solvated in an octahedral TIP3P water box with a minimum 5 Å buffer from the protein surface and neutralised with Na⁺ and Cl⁻ ions, adjusted to a final concentration of 150 mM. The fully assembled complexes measured approximately 17 × 17 × 17 nm³ and contained ∼400,000 atoms.

All systems underwent 2,000 steps of steepest-descent energy minimisation to remove steric clashes. The systems were then gradually heated to 300 K over 1 ns with a 2fs time step, applying harmonic restraints (1 kcal·mol⁻¹·Å⁻²) on all non-hydrogen atoms. Subsequent equilibration was performed for 20 ns at 1 atm and 300 K to achieve the correct system density. Following equilibration, all restraints were removed, and production MD simulations were performed for a minimum of 2 μs under the NPT ensemble (constant number of particles, pressure = 1 bar, temperature = 300 K). Trajectories were analysed to assess conformational behaviour, structural arrangement, and intermolecular interactions across wild-type and L8A systems.

## Acknowledgements

P.M. acknowledges funding support from SERB (ECR/2017/003431), UoH-IoE-RC5-22-011, MoE-STARS/STARS-1/PID (STARS1/634), HGK-IYBA (BT/11/IYBA/2018/08) and DBT BUILDER grant (BT/INF/22/SP41176/2020). P.M. is a recipient of long term ICMR-DHR Fellowship from ICMR. PM duly acknowledges the support of Ramalingaswami Fellowship, Department of Biotechnology, GOI. AAP is a recipient of PMRF, GOI. SS is a recipient of CSIR-SRF, GOI.

## Supplementary Figure Legends

**Figure S1.**
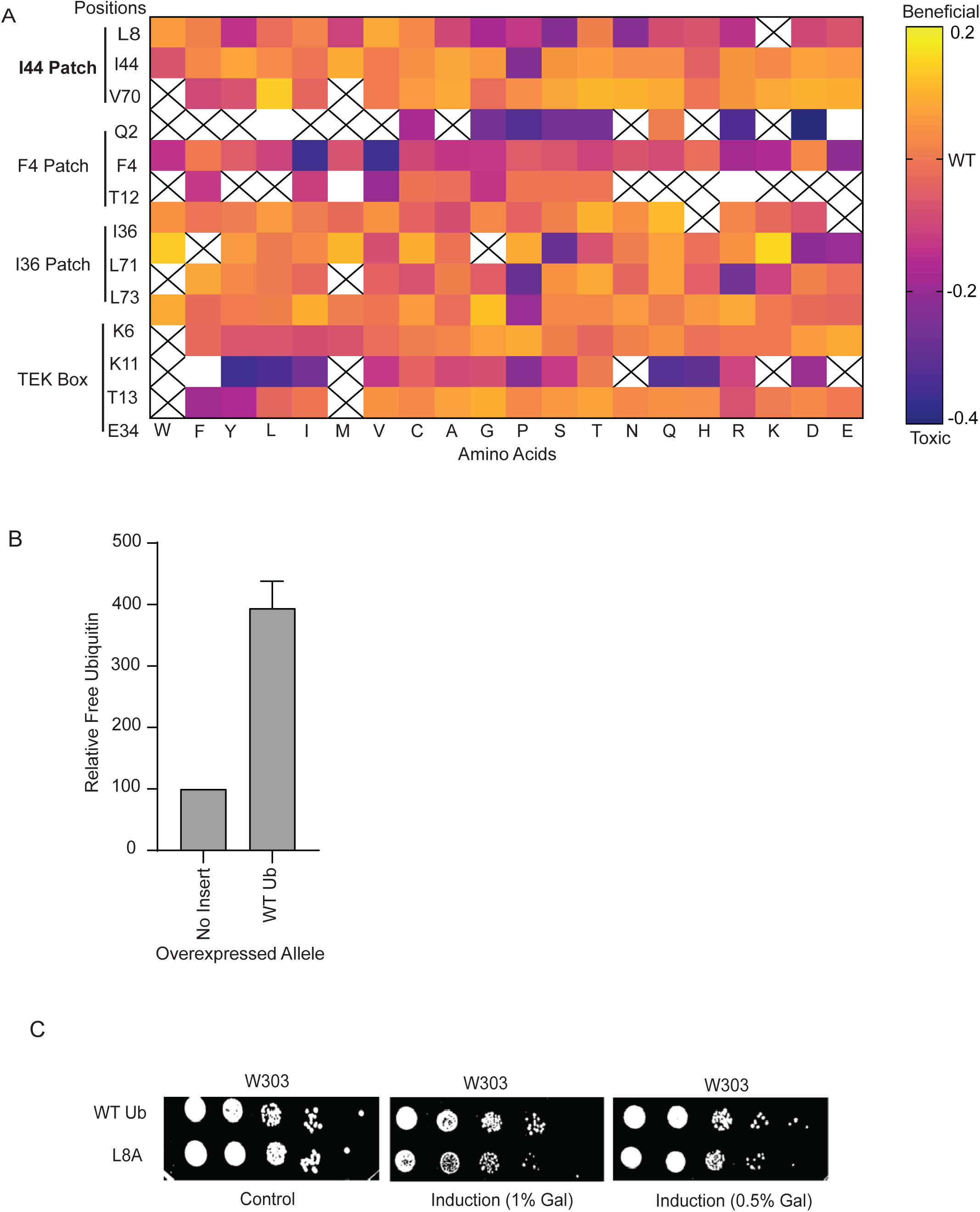
(A). Heat map showing growth phenotypes of ubiquitin mutants targeting different hydrophobic patches (data from our previous study). Overexpression of these mutants results in beneficial (yellow) or deleterious (purple) growth phenotypes. (B). Quantification of the increase in free ubiquitin levels upon overexpression of WT Ub and L8A in W303 cells. Data are presented as mean ± SEM (n = 3). (C). Growth analysis of yeast cells overexpressing WT Ub or L8A along with an endogenous copy of WT Ub, induced for 3 hours in media containing 2% or 1% galactose.

**Figure S2.**
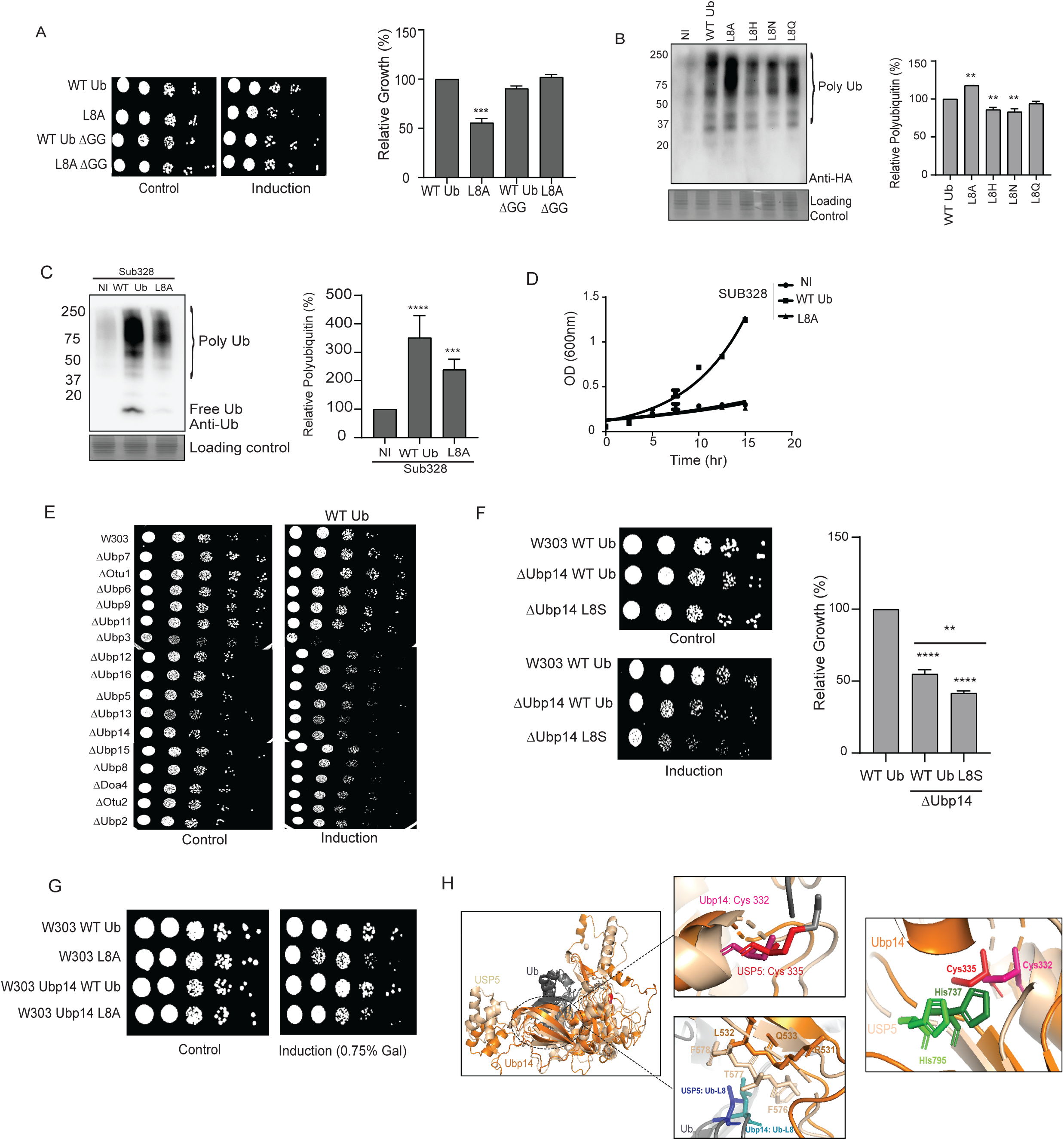
(A). Growth analysis of W303 cells on solid media overexpressing ΔGG mutant in L8A or WT sequence. Spot dilution assays were performed on tryptophan-deficient synthetic media, with dextrose as the control medium and galactose for mutant induction. The bar graph shows growth rates of ubiquitin variants relative to overexpressed WT Ub in cells grown on solid medium for 2–3 days. Data are presented as mean ± SEM (n = 2 independent experiments). Statistical significance was determined using one-way ANOVA with Dunnett’s post hoc test comparing each mutant with wild type. (B). Western blot image for specific accumulation of HA tagged polyubiquitinated substrates in cells overexpressing the L8 mutants alongside endogenous WT Ub background in W303 cells. Data are presented as mean ± SEM (n = 3 independent experiments). Statistical significance was determined using one-way ANOVA with Dunnett’s post hoc test comparing each mutant with W303 wild type. (C). Western blot image for SUB328 strain overexpressing WT Ub and L8 mutant. Data are presented as mean ± SEM (n = 3 independent experiments). Statistical significance was determined using one-way ANOVA with Dunnett’s post hoc test comparing each mutant with SUB328 wild type. (D). Growth of Sub328 cells in liquid media with the strain overexpressing either the WT Ub or the mutant Ub followed for 15 hours post induction. The cells were diluted to mid-log phase of growth. (E). Growth of all the non essential DUBs strains without (control) and with (induction) overexpressing the WT Ub in synthetic growth media deficient for Ura. (F). Growth analysis of yeast cells overexpressing L8S mutant alongside an endogenous copy of WT Ub. Spot dilution assays were performed on tryptophan-deficient synthetic media, with dextrose as the control medium and galactose for mutant induction. The bar graph shows growth rates of ubiquitin variants relative to overexpressed WT Ub in cells grown on solid medium for 2–3 days. Data are presented as mean ± SEM (n = 2 independent experiments). Statistical significance was determined using one-way ANOVA with Dunnett’s post hoc test comparing each mutant with wild type. (G). Growth analysis of yeast cells overexpressing Ubp14 induced with copper for 9 hours and WT Ub or L8A induced with shift to 1% or 0.75% galactose media for 3 hours. (H). Structural alignment of the catalytic domain of Ubp14 (AlphaFold model, shown in orange) and USP5 (PDB: 3IHP, modified, shown in wheat) bound to ubiquitin. The overlapping ubiquitin-binding region is indicated with a dotted circle. The upper right zoomed panel shows the active cysteine residues of USP5 (red) and Ubp14 (pink) represented as sticks. The lower right zoomed panel highlights residues of USP5 and Ubp14 in proximity to ubiquitin L8. The right panel shows the structural alignment of the Cys box and His box motifs of USP5 and Ubp14.

